# STAT5b is a key effector of NRG-1/ERBB4-mediated cardiomyocyte growth

**DOI:** 10.1101/2022.10.05.510958

**Authors:** Katri Vaparanta, Anne Jokilammi, Ilkka Paatero, Johannes A. Merilahti, Juho Heliste, Karthik Amudhala Hemanthakumar, Riikka Kivelä, Kari Alitalo, Pekka Taimen, Klaus Elenius

## Abstract

The growth factor neuregulin-1 (NRG-1) regulates hypertrophic and hyperplastic myocardial growth and is currently under clinical investigation as a treatment for heart failure. We have previously demonstrated that an isoform of the NRG-1 receptor ERBB4 (ERBB4 JM-b) expressed in cardiomyocytes selectively regulates the activation of STAT5b. To explore the role of STAT5b in NRG-1/EBBB4 mediated cardiomyocyte growth, several *in vitro* and *in vivo* models were utilized. The downregulation of NRG-1/ERBB4 signaling consistently reduced STAT5b activation and transcription of STAT5b target genes *Igf1, Myc* and *Cdkn1a* in murine *in vitro* and *in vivo* models of myocardial growth. *Stat5b* knock-down in primary cardiomyocytes ablated NRG-1-induced cardiomyocyte hypertrophy. Stat5b was activated during NRG-1-induced hyperplastic myocardial growth and chemical inhibition of the Nrg-1/Erbb4 pathway led to the loss of myocardial growth and Stat5 activation in zebrafish embryos. Moreover, CRISPR/Cas9-mediated knock-down of *stat5b* in zebrafish embryos resulted in reduced myocardial growth and heart failure as indicated by reduced ventricular ejection fraction. Dynamin-2 was discovered to control the cell surface localization of ERBB4 and the chemical inhibition of dynamin-2 downregulated NRG-1/ERBB4/STAT5b signaling in models of hypertrophic and hyperplastic myocardial growth. Finally, the activation of the NRG-1/ERBB4/STAT5b signaling pathway was explored in clinical samples representing pathological cardiac hypertrophy. The NRG-1/ERBB4/STAT5b signaling pathway was differentially regulated both at the mRNA and protein levels in the myocardium of patients with pathological cardiac hypertrophy as compared to myocardium of control subjects. These results establish the role for STAT5b, and dynamin-2 in NRG-1/ERBB4-mediated myocardial growth.

## Introduction

Neuregulin-1 (NRG-1) has been implicated as a central regulator of both hypertrophic and hyperplastic cardiomyocyte growth ^1,2^. NRG-1 activates ERBB receptor tyrosine kinases by inducing their dimerization and subsequential transphosphorylation, and is known to signal through the ERBB2/ERBB4 heterodimer in the myocardium ^3–5^. Of these two ERBB receptors, only ERBB4 can directly bind NRG-1. ERBB2 is activated by dimerizing with the NRG-1-bound ERBB4 receptor^6^. Genetic knock-out or chemical inhibition of NRG-1, ERBB2 or ERBB4 in zebrafish and mouse models has been shown to lead to a distinct heart phenotype during embryogenesis characterized by dilated ventricles, thin ventricular walls and loss of trabeculae ^3,5,7,8^, and to dilated cardiomyopathy in the adulthood ^9–11^. After successful phase II trials ^12,13^, NRG-1 is currently tested in phase III clinical trials as a treatment for heart failure^14^.

The NRG-1/ERBB pathway has been shown to activate PI3K/AKT, MAPK, and SRC/FAK pathways in cardiomyocytes ^2,15–19^, none of which has been demonstrated to directly control NRG-1/ERBB-mediated cardiomyocyte growth in *in vitro* or *in vivo* settings. Instead, the PI3K/AKT pathway has been affiliated with cardiomyocyte survival, DNA synthesis and glucose uptake ^1,16–18^, the MAPK pathway with protein synthesis and sarcomere organization ^2^, and the SRC/FAK pathway with cytoskeletal remodeling ^15^.

In our previous study we discovered that the transcription factor STAT5b is selectively activated by the JM-b isoform of ERBB4 ^20^, the ERBB4 isoform preferentially expressed by cardiomyocytes (Supplementary Figure 1A) ^21,22^. Of the two *STAT5* genes, *STAT5b* is also the major *STAT5* gene expressed in the heart (Supplementary Figure 1B). Since chemical inhibition of STAT5 has previously been shown to attenuate myocardial growth induced by angiotensin II ^23^ or by pressure overload in mice, we set out to explore the role of STAT5b in NRG1/ERBB4-mediated myocardial growth. The analyses validated in both *in vitro* and *in vivo* models demonstrated that the NRG-1 receptor ERBB4, the ERBB4-activated intracellular signaling molecule STAT5b, and the ERBB4 trafficking-regulator dynamin-2 mediate both hypertrophic and hyperplastic NRG-1-induced cardiomyocyte growth. In addition, we discovered altered signaling of the NRG-1/ERBB4/STAT5b/dynamin-2 pathway in the myocardia of patients suffering from pathological cardiac hypertrophy.

**Figure 1.**
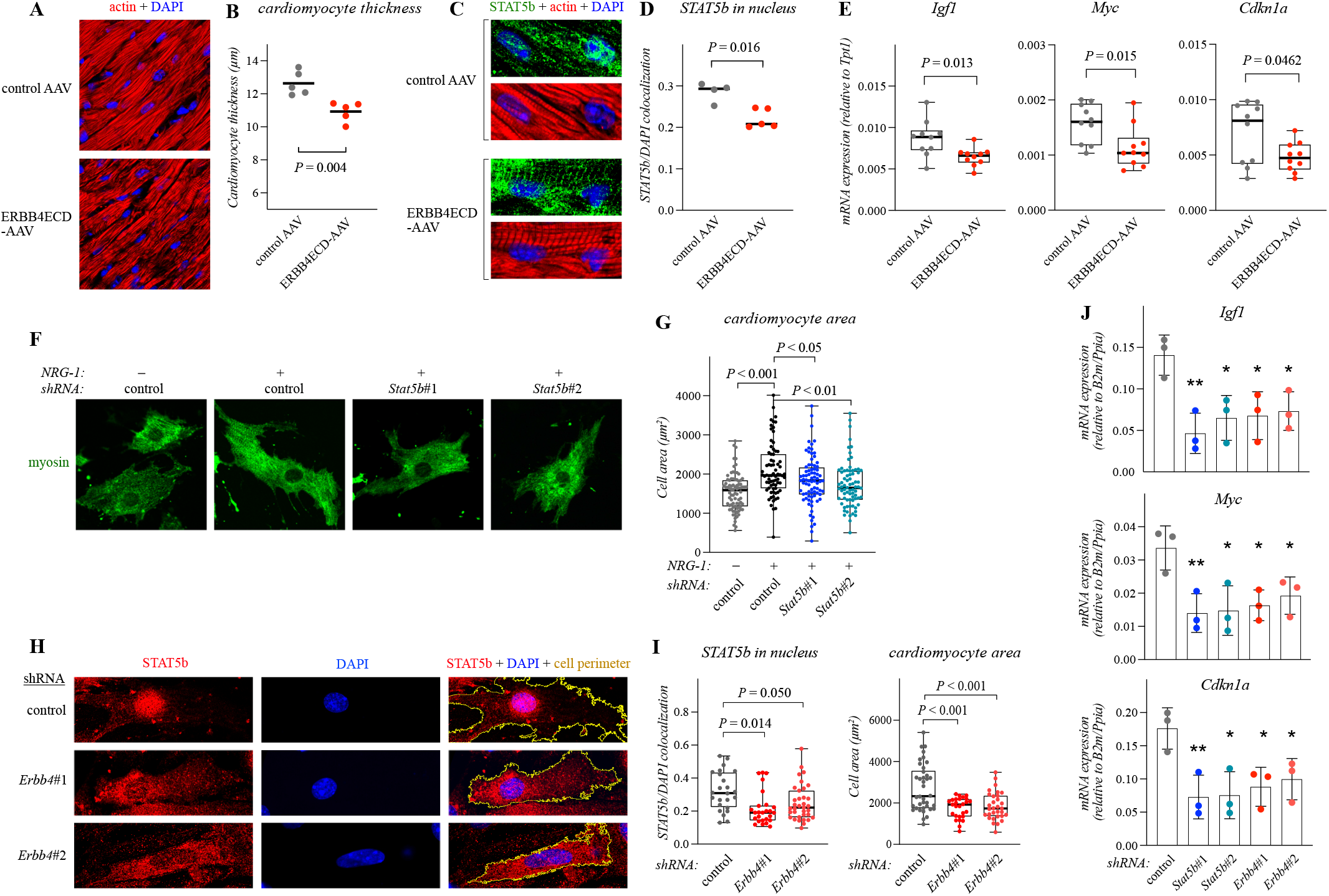
NRG-1 promotes cardiomyocyte hypertrophy via ERBB4 and STAT5b. **A-B**: Confocal images (A) and quantification (B) of immunofluorescence staining of actin filaments (phalloidin) and nuclei (DAPI) in cardiomyocytes. Images of cryosections from hearts of mice transduced with either control (control AAV) or ERBB4 ectodomain-encoding (ERBB4ECD-AAV) adeno-associated viruses are shown. One dot in the scatterplot corresponds to the average cardiomyocyte thickness as defined by the thickness of the myofibril bundle per cardiomyocyte in 3-5 randomly acquired images of stained sections from one heart (n = 5). Two-tailed unpaired T-test was used for statistics. **C-D:** Confocal images (C) and quantification (D) of immunofluorescence staining of STAT5b, actin filaments (phalloidin) and nuclei (DAPI) in cardiomyocytes of cryosections from the hearts of mice transduced with either control- or ERBB4ECD-AAV. PanelD depicts quantification of colocalization of STAT5b- and DAPI-specific signals. One dot in the boxplot corresponds to the median value in 20-50 images of stained sections from one heart (n = 4-5). Two-tailed unpaired T-test was used. **E:** Real-time RT-PCR analysis of expression of the indicated STAT5b target genes in the heart of mice transduced with either control- or ERBB4ECD-AAV. One dot corresponds to analysis of one heart (n = 10; combined from two replicate experiments). Two-tailed unpaired T-test was used. **F-G:** Confocal images (F) and quantification (G) of immunofluorescence staining of myosin heavy chain in primary mouse cardiomyocytes. The cells were treated with either control or *Stat5b*-targeting shRNAs and stimulated with NRG-1 for two days. The myosin immunoreactivity was used to quantify cell area. One dot in the boxplot corresponds to one cell (n = 73-83; combined from four replicate experiments). Non-parametric Kruskal-Wallis ANOVA was used. Post hoc analyses were conducted with the Mann Whitney U-test and the resulting P-values corrected with the method of Benjamini, Krieger and Yekutieli. **H-I:** Confocal images (H) and quantification (I) of immunofluorescence staining of STAT5b and nuclei (DAPI) in mouse cardiomyocytes. The cells were treated with either control or *Erbb4*-targeting shRNAs. Panel I depicts quantification of colocalization of STAT5b- and DAPI-specific signals (left) and the cell perimeter (right). One dot in the boxplots corresponds to one cell (n = 24-35; combined from two replicate experiments). Non-parametric Kruskal-Wallis ANOVA and Dunn’s multicomparison test (colocalization analysis) and Brown-Forsythe one-way ANOVA and Dunnett’s multicomparison test (cardiomyocyte size analysis) with multiple test correction were used. **J:** Real-time RT-PCR analysis of expression of the indicated STAT5b target genes in mouse cardiomyocytes. The cells were treated with either control, *Stat5b*- or *Erbb4*-targeting shRNAs and stimulated with NRG-1 for 30 minutes. One dot corresponds to the mean of technical repeats in one experiment and the whiskers the standard deviation (n = 3; replicate experiments). One-way ANOVA and the Dunnett’s multicomparison test with multiple test correction was used. *, *P* < 0.05; ** *P* < 0.01; ***, *P* < 0.001.

## Materials and Methods

### Resources table

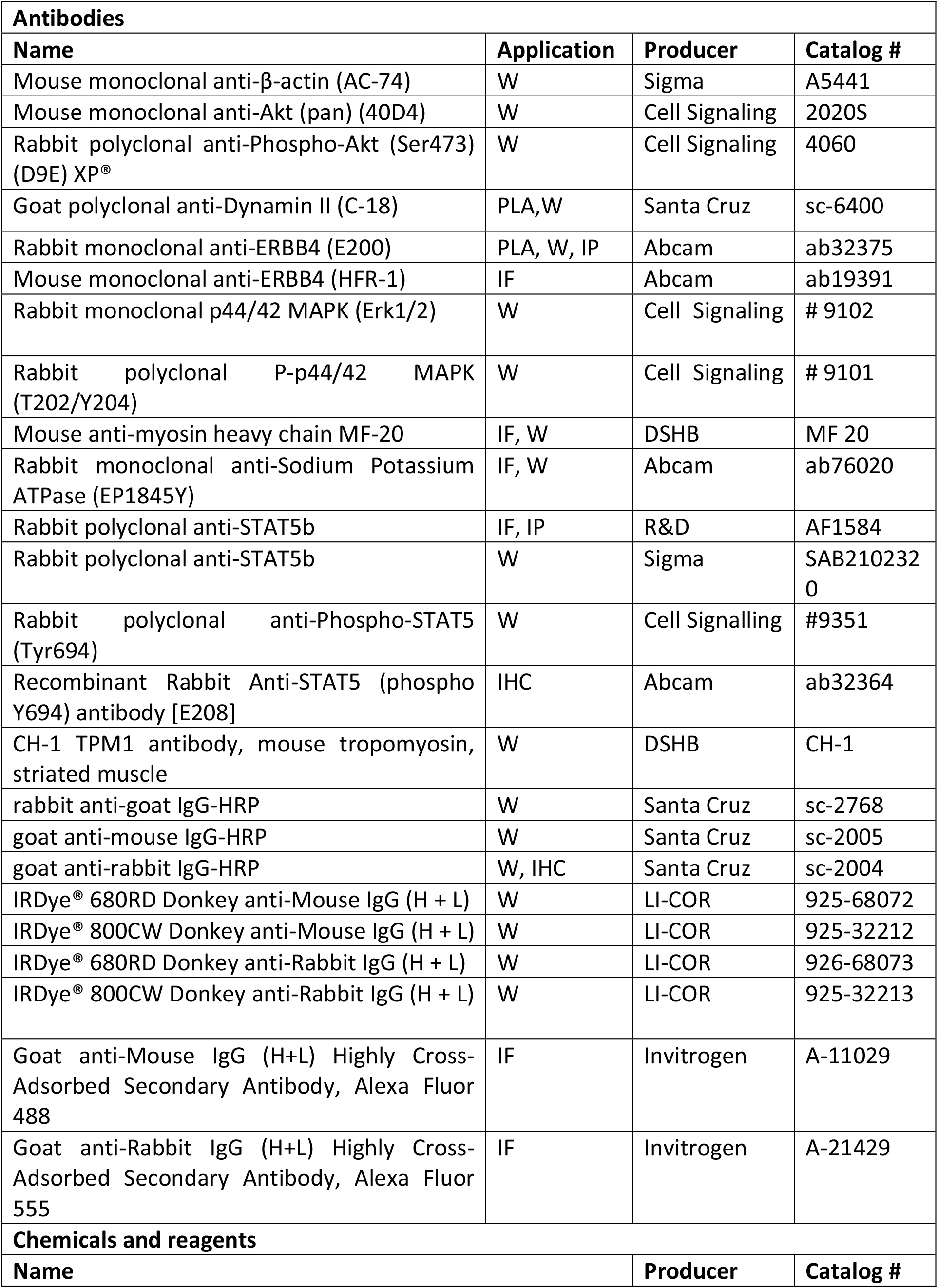
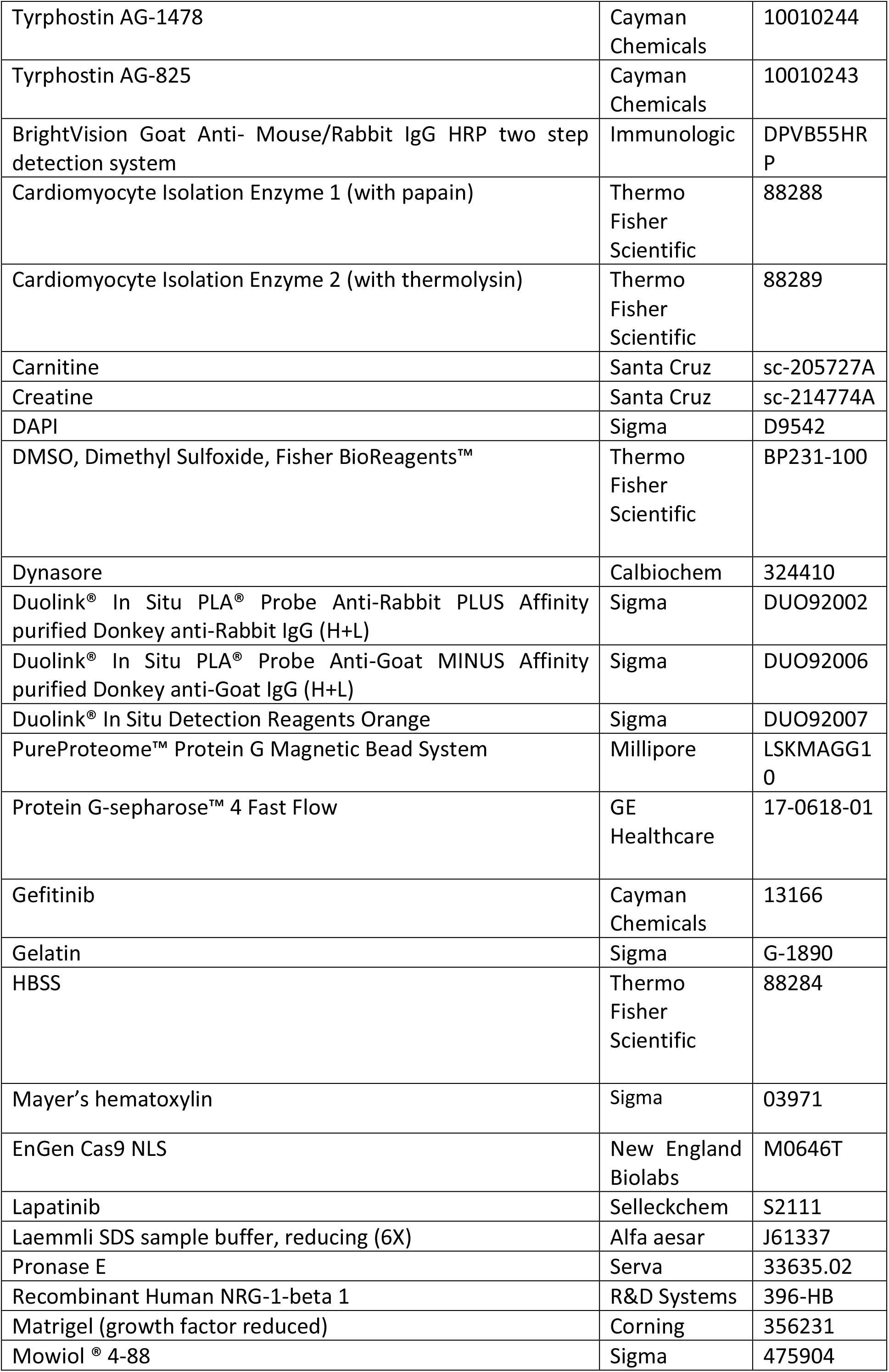
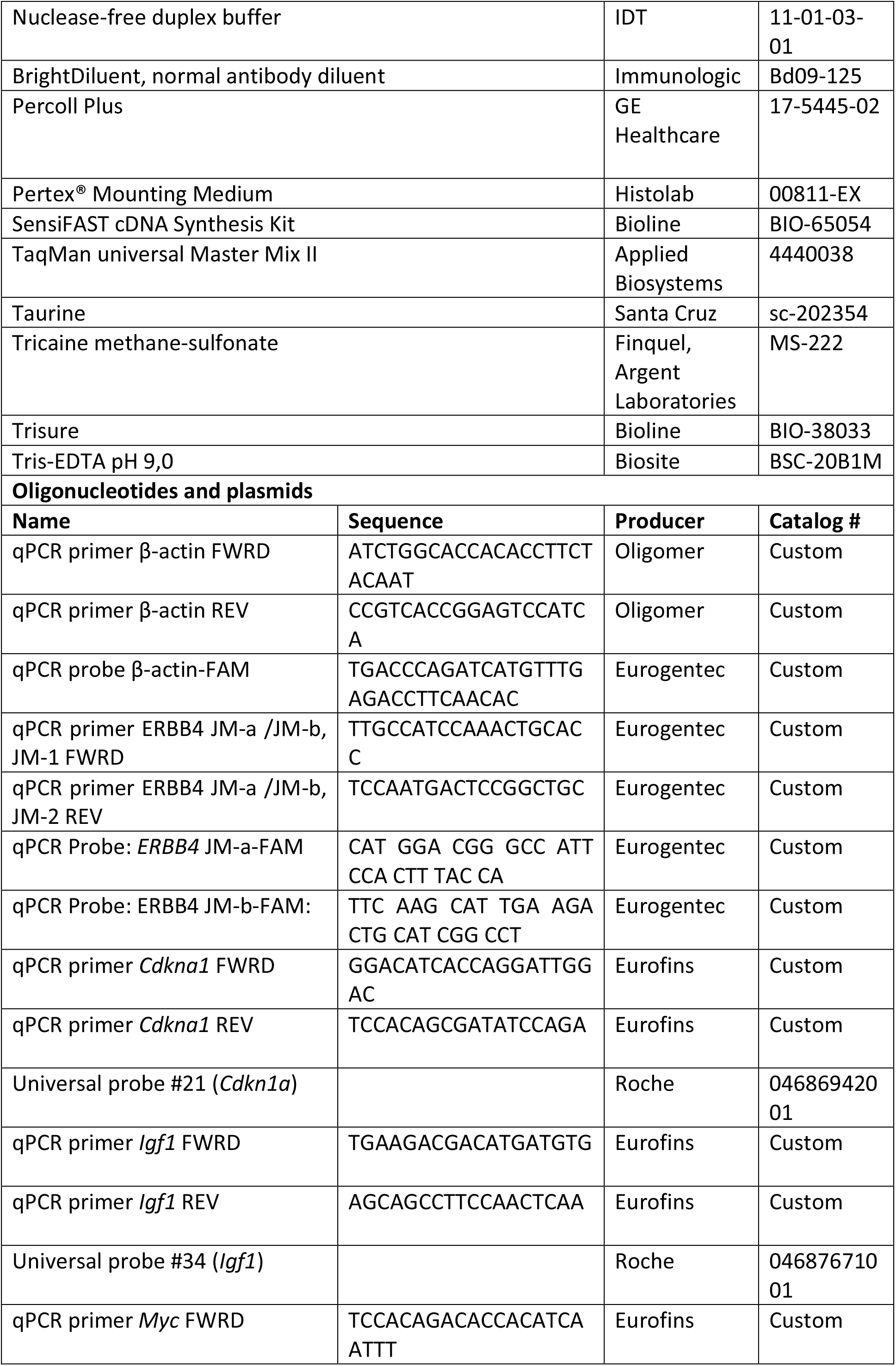
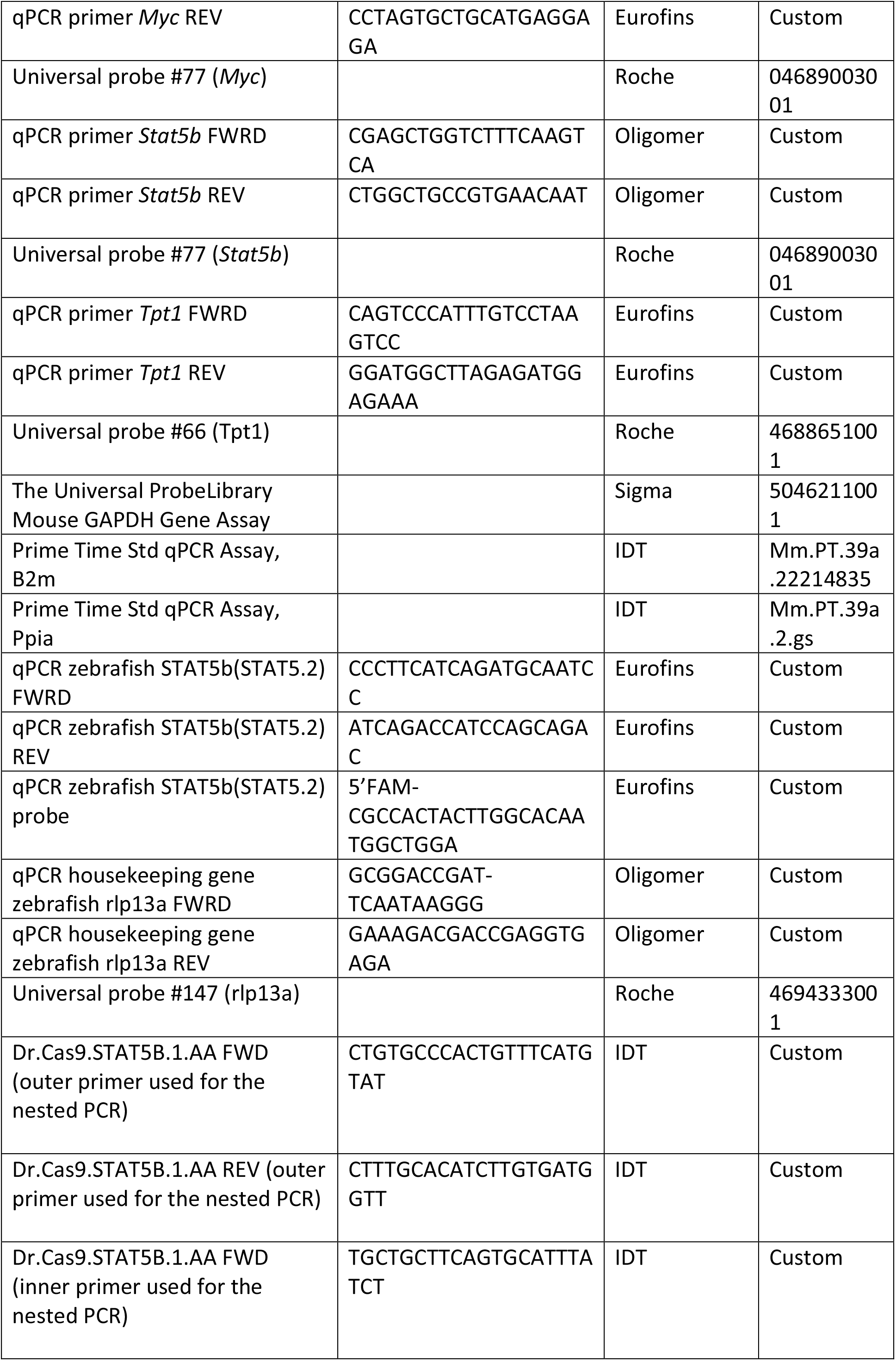
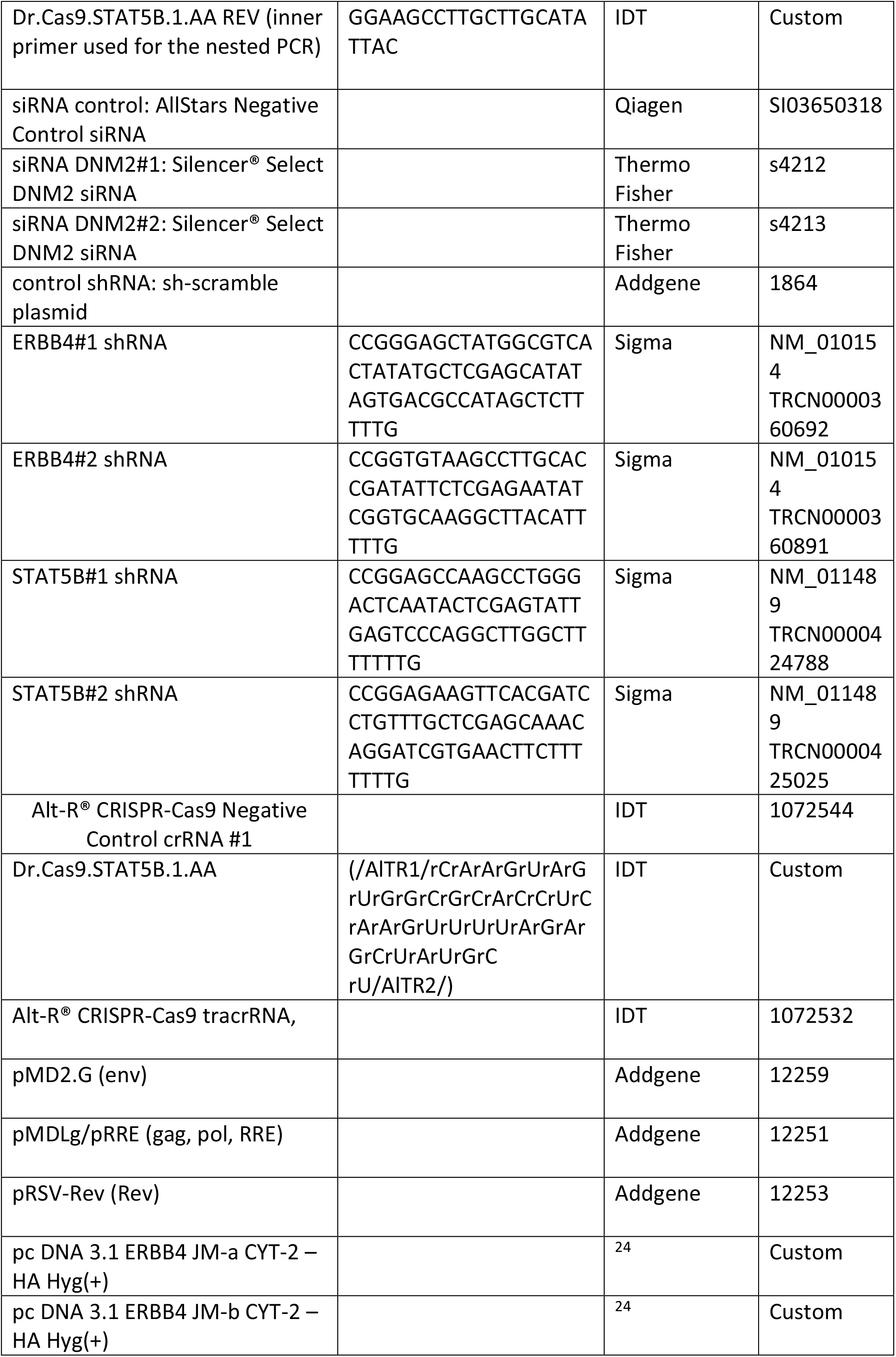
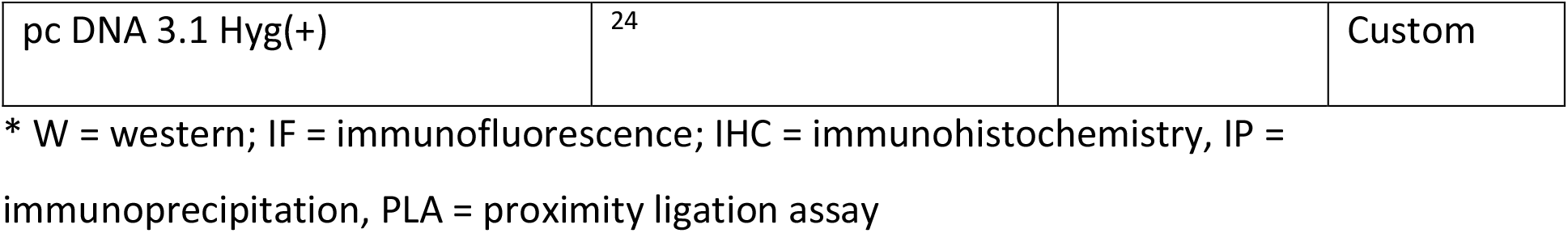

### Isolation and culture of primary murine neonatal cardiomyocytes

NMRI male mice (Janvier Labs) were housed in the Central Animal Laboratory of University of Turku. The housing, upkeep and euthanasia of animals were conducted according to Directive 2010/63/EU and under license from The Ministry of Agriculture and Forestry of Finland. The neonatal mice were sacrificed by decapitation. Freshly dissected neonatal (≤ 3 days post partum) hearts were minced and washed twice with ice-cold HBSS (Hank’s balanced balt solution). The hearts were enzymatically digested with Cardiomyocyte Isolation Enzymes 1 and 2, papain, and thermolysin, according to manufacturer’s instructions, except that thermolysin was used 1:5 of its suggested volume. The samples were washed twice with ice cold ADS buffer ^25^, mechanically agitated, and centrifuged at 900 g for 6 minutes in ADS buffer. The isolated cells were purified with a Percoll gradient, as previously described ^25^. Cell concentration and viability of cardiomyocytes was determined with TC20 Automated Cell Counter (Bio-Rad). Cells were plated on 1% gelatin-coated plates and maintained in plating medium [68% DMEM (Lonza), 17% 199 (Sigma), 10% horse serum (PromoCell), 5% fetal calf serum (Biowest), pH 7.2 ^25^]. After two days, the medium was changed to serum-free maintenance medium [80% DMEM, 20% 199, pH 7.2, 5 mM Creatine, 2 mM Carnitine, 5 mM Taurine ^25^]. For immunofluorescence analyses, the primary neonatal cardiomyocytes were plated in coverslips coated with 2% Matrigel and cultured in plating medium.

### Zebrafish husbandry

Zebrafish (*Danio rerio*) of casper strain (*roy, mitfa*) ^26^ were housed at Zebrafish Core of Turku Bioscience Center, University of Turku and Åbo Akademi according to standard procedures ^27^. Embryos were obtained through natural spawning in dedicated breeding tanks. After collection, the embryos were flushed and incubated in E3-medium at +28.5°C in a humidified incubator. The zebrafish embryos were anesthetized by administrating single dose of tricaine (Ethyl 3-aminobenzoate methanesulfonate) into the culture medium at end concentration of 200mg/l. The zebrafish embryos were euthanized under terminal anesthesia by fixation in 4% paraformaldehyde or direct homogenization using motorized pellet pestle (model Z359971, Sigma-Aldrich). The zebrafish embryos at the age (< 5dpf) used in experiments are not considered as protected animals as defined by Directive 2010/63/EU. The adult zebrafish were housed and used in matings according to Directive 2010/63/EU and under license MMM/465/712-93 (Ministry of Forestry and Agriculture).

### Patient sample characteristics

The formalin-fixed paraffin-embedded myocardial tissue sections were acquired from Auria biobank though an application process (application number: AB20-3971). The Finnish Biobank Act (Finlex 688/2012) enables the use of biobank resources in research. The patients whose samples have been used in the study have given their consent to use their samples in research to Auria biobank. The research conforms to the principles outlined in the Declaration of Helsinki. The cohorts included myocardial samples obtained from 19 patients who underwent medical autopsy at Turku University Hospital between 2008–2017. The hypertrophy cohort (median age 67; 23% female) included 6 samples from patients with aortic valve stenosis, 4 samples from patients with alcoholic cardiomyopathy and 3 from patients with idiopathic cardiomyopathy. The control cohort (median age 48; 50% female) included 6 cardiac samples taken from autopsied patients with non-cardiac diseases (2 cases of cirrhosis, 1 case of metastatic pancreatic carcinoid, 1 case of pulmonary embolism, 1 case of sepsis due to Streptococcus pneumoniae and 1 case of cholangiocarcinoma). All the cases in the control cohort were negative for cardiomyocyte hypertrophy based on histopathological analysis.

### Adeno-associated virus-treated mouse samples

The adeno-associated virus (AAV)-treated mouse samples were acquired from the experiments described in Kivelä et al., 2019 ^28^. Before tissue collection, mice were injected intraperitoneally with 80mg/kg of ketamine and 10mg/kg of xylazine and euthanized by cervical dislocation. The local committee appointed by the district of southern Finland approved the animal experiments. FELASA (Federation of European Laboratory Animal Science Associations) guidelines and recommendations were followed. All the animal procedures and experiments were performed under the EU directive 2010/63/EU guidelines of the European parliament on the protection of animals used for scientific purposes.

### Viral transduction

Lentiviral vectors were produced in HEK293T cells using a third generation lentiviral packaging system ^29^. The HEK293T cells were cultured in DMEM supplemented with 10% FBS, 5 mM Glutamine, and 1 U of penicillin and streptomycin. Cells were co-transfected overnight with pRSV-Rev, pMDLg/pRRE, pMD2.G, and pLKO.1-puromycin encoding one of the shRNA constructs. Medium was changed 24 hours after transfection, and virus-containing medium was collected at 48 and 72 hours and filtered. Primary cardiomyocytes were infected with the lentiviruses at a multiplicity of infection (MOI) of 1 in the presence of 8 μg/ml polybrene. The medium was changed to serum-free maintenance medium [80% DMEM, 20% 199, pH 7.2, 5 mM Creatine, 2 mM Carnitine, 5 mM Taurine (Louch et al., 2011)] after 5 hours. The cells were analyzed 24 hours after infection.

### Neuregulin stimulation and chemical inhibition

For western analyses, cells were stimulated with 50 ng/ml of human recombinant NRG-1β for 2 minutes where indicated. For real-time RT-PCR analyses, the cells were incubated with 50 ng/ml NRG-1β for 30 minutes. For immunofluorescence analyses of hypertrophic growth, the isolated cardiomyocytes were incubated with 50 ng/ml NRG-1β for 2 days. To inhibit signaling, isolated cardiomyocytes were treated with the ERBB inhibitors 10 µM AG 1478, 10 µM AG 825 for 1 hour or 50 µM dynasore for 3-6 hours. To analyze the effect of ERBB or dynamin inhibition to the morphology and function of the heart of embryonic zebrafish, zebrafish embryos were treated with 10 µM AG 1478, 30 µM lapatinib, 30 µM gefitnib or 50 µM dynasore at 2 dpf for 2-3 days. To analyze the effect of ERBB or dynamin inhibition on downstream signaling with western analyses, zebrafish embryos at 4-5 dpf were treated with 10 µM AG 1478, 30 µM lapatinib, 30 µM gefitinib or 50 µM dynasore for 1-3 hours. To analyze the effect of NRG-1 on the morphology and function of the heart of embryonic zebrafish, 100 pg of recombinant human NRG-1β or control substance (0.2% BSA in PBS) was microinjected to the pericardial sac of 2 dpf zebrafish embryos and the embryos were analyzed at 4-5 dpf. To analyze the effect of NRG-1 to downstream signaling with western analysis, 100 pg of recombinant human NRG-1β or control substance with 100 mM Na_3_VO_4_ was microinjected to the pericardial sac of 4-5 dpf zebrafish embryos and lysed after 20 minutes.

### Immunofluorescence and immunohistochemistry

For immunofluorescence analysis of isolated cardiomyocytes, samples were fixed and permeabilized with methanol for 10-15 minutes at -20 ⁰C and stained with the indicated primary antibodies. To remove non-specific binding, 3% BSA in PBS was used as a blocking agent. Primary antibodies were detected with Alexa-conjugated secondary antibodies and nuclei were visualized with DAPI. Mowiol was used for mounting. The samples were imaged with Zeiss LSM 780 confocal microscope with the 40x and 63x Zeiss C-Apochromat objectives (Numerical Aperture: 1.2).

For immunofluorescence analysis of murine cardiac cryosections, the cryosections were allowed to reach room temperature and were rehydrated in PBS for 30 minutes. Sections were blocked by 3% BSA in PBS, and stained with anti-STAT5b (1:20; R&D) for 3 hours. The sections were washed 3 times with PBS and incubated with the secondary Alexa Fluor 488 goat anti-rabbit antibody. Actin and nuclei were visualized with Alexa Fluor 555-conjugated phalloidin and DAPI (4′,6-diamidino-2-phenylindole), respectively. Coverslips were mounted with Mowiol. The samples were imaged with Zeiss LSM 780 and 880 confocal microscopes with the 40x Zeiss C-Apochromat and the 40x Zeiss LD LCI Plan-Apochromat objectives (Numerical Aperture: 1.2).

For immunofluorescence analysis of zebrafish embryos, the embryos were fixed and permeabilized with 4% PFA and 0.2% Triton X-100 in PBS for 30 minutes. The embryos were washed 4 times with 0.2% Tween 20 in PBS and blocked with 3% BSA and 0.2% Tween 20 in PBS for 1 hour. The primary and Alexa 555- or 488-conjugated secondary antibodies were incubated overnight in +4⁰C under gentle agitation. The embryos were washed 4 times with 0.2% Tween 20 in PBS for 30 minutes under gentle agitation after primary and secondary antibody incubation. The stained embryos were imaged with Zeiss LSM 780 confocal microscope with the 40x Zeiss C-Apochromat objective.

Human heart tissue section were analyzed by immunohistochemistry in the Histology Core Facility of the Institute of Biomedicine, University of Turku. After deparaffinization and rehydration, the sections were incubated in Tris-EDTA pH 9.0 in microwave oven for 7 minutes with 600 W followed by 7 minutes with 450 W, for antigen retrieval. The sections were incubated in the presence of hydrogen peroxide to inhibit endogenous enzyme function and normal antibody diluent to prevent non-specific binding of proteins. The sections were stained with an automated Labvision autostainer (Thermo-Fisher Scietific) with anti-pSTAT5 (1:1000) for 60 minutes in room temperature. Primary antibody was detected with BrightVision Goat Anti-Mouse/Rabbit IgG HRP two step detection system with horseradish peroxidase (HRP)-conjugated secondary antibodies. Slides were counterstained with Mayer’s hematoxylin for 1 minute at room temperature, dehydrated and mounted with Pertex (Histo-Lab). The stained section were imaged with Panoramic P1000 slide scanner (3DHISTECH).

### Proximity ligation assay

The proximity ligation assay of dynamin-2/ERBB4 association was performed according to manufacturer’s protocol with anti-mouse minus and anti-goat plus probes. A 1:100 dilution of primary dynamin-2 and ERBB4 antibodies was used. The primary ERBB4 antibody was omitted from the technical control.

### Western analysis

The zebrafish embryos were ground with an electronic pestle into 6x Laemmli buffer and incubated at 100 ⁰C for 10 minutes. The samples were centrifuged and loaded on to SDS-PAGE gels, run and transferred to nitrocellulose membranes. The membranes were probed with indicated primary antibodies and IR-conjugated secondary antibodies and imaged with the Li-COR Odyssey system. The densitometric quantitation was performed with Image Studio Lite.

### RNA extraction and real-time RT-PCR

Total RNA was extracted using TRIsure RNA isolation reagent according to manufacturer’s instructions. RNA concentrations were determined using a Nanodrop spectrophotometer (Thermo Fisher Scientific), and complementary DNA was synthesized with Sensifast cDNA Synthesis Kit from 1 µg of total RNA. Real-time RT-PCR analyses of gene expression were performed with 500 nM of primers, 100 nM of 5’ 6-FAM-labeled probe and TaqMan universal Master Mix II according to manufacturer’s protocol. Thermal cycling was performed with QuantStudio 12K Flex Real-Time PCR System (Thermo Fisher Scientific). The relative real-time RT PCR values were determined by normalizing the Ct values of the gene of interest with the Ct values of the housekeeping genes as follows: 2^Ct_gene_of_interest – 2^Ct_housekeeping_gene. The Ct values of the genes of interest in isolated cardiomyocytes were normalized to the average Ct values of *B2m* and *Ppia* or *Gapdh* and *Actin* housekeeping genes. The Ct values of the genes of interest in cardiac tissue samples were normalized to the Ct value of the *Tpt1* gene.

### *In vivo* imaging of zebrafish embryos

The embryos were anaesthetized with Tricaine methane-sulfonate and molded into low-melting point agarose. The fast time laps videos of embryos were taken with HC Image Software of Zeiss AxioZOOM.V16 with 1.0x PlanApo Z objective with 80 x magnification.

### CRISPR/Cas9 mediated knock-down of STAT5b in zebrafish embryos

The CRISPR/Cas9 mediated knock-down was carried out largely as described in ^30^. Guide RNAs were generated by combining 5 µl of crRNAs (100 µM) and tracRNA (100 µM). Duplexes were annealed by heating to 95 °C for 5 minutes, cooled down at 0.1 °C/second to 25 °C, and incubated at 25 °C for 5 minutes. Ten µl of nuclease-free duplex buffer (IDT) was added to reach a final concentration of 25 µM and duplexes were stored stored at -20 °C.

To generate crRNA:tracRNA:Cas9 complexes, 1 µl of duplex, 1.25µl of EnGen Cas9 NLS solution (NEB), 1.75 µl H_2_O, and 1 µl of phenol red solution (Sigma) were mixed together and incubated for 5 minutes at 37 °C prior to injection. Next, 2.3 nl of solutions was injected into 1-4 cell stage zebrafish embryos of casper line using Nanoject II microinjector (Drummond Scientific). After injection, the embryos were placed to a +28.5 °C incubator in E3-medium supplemented with 1:100 Pen-Strep. At 1 dpf, dead embryos were removed and 15 µl of pronase E was added to facilitate hatching. The embryos were analyzed at 4 dpf.

### Image analysis

The area of isolated cardiomyocytes and zebrafish embryo ventricles, and the thickness of ventricles was measured with Fiji v1.53c. The ejection fraction of zebrafish ventricle was determined by measuring the average width of the ventricle at systole and diastole with Fiji and fitting the values to the simplified Quinones equation: (EDD^2 - ESD^2) / EDD^2 * 100% + 10%. The ventricular width, measured from outer margins of the myocardium, at diastole and at systole was used as the end-diastolic dimension (EDD) and end-systolic dimension (ESD), respectively. The amount of PLA signals was measured with the particle analyzer function of Fiji and the location of PLA signals was analyzed with SpatTrack v2 ^31^. The colocalization of immunofluorescence signals was measured with the algorithm of Villalta et al ^32^. Four colocalization measures (Pearson correlation, Manders overlap, M1/M2, k1/k2) were produced of which k1/k2 (*k*1 = ∑(*S*_1_ × *S*_2_)/ ∑(*S*_1_) and *k*2 = ∑(*S*_1_ × *S*_2_)/ ∑(*S*_2_), where S_1_ and S_2_ are the intensity of the signal from channel 1 and 2, respectively) is visualized in the images. The relative intensity of the DAB staining was measured from color deconvoluted images with Fiji. The average intensity in the cardiomyocyte nuclei to cardiomyocyte cytosol (background) was used. The intensity of immunofluorescence signal was measured with Fiji.

### Data and statistical analyses

Matlab R2016a and Graph Pad Prism 8.4.2 were used for statistical analysis. The normality and homoscedasticity of each dataset was tested with Shapiro-Wilkis, Kolmogorov-Smirnov, Bartlett’s and Brown-Forsythe test and parametric, parametric testing with variance correction or non-parametric testing was chosen accordingly. All data combined from independent experiments was median-normalized except for the data in Figure 1J. The data in 1J were normalized to the range to fit the combined data into a normal distribution. The transcriptome datasets of subjects with pathological hypertrophy and control subjects were downloaded from the Gene expression omnibus database ^33,34^. The principle component analysis of the pre-existing datasets was performed with R version 3.6.1. The statistical significance of the categorization of the dimensionality-reduced values was calculated by estimating an empirical probability distribution for the relative distance and drawing the cumulative probability from the corresponding cumulative probability function. The relative distance was determined by the sum of all Euclidian distances within the category divided by the sum of all Euclidian distances between categories. The empirical probability distribution was simulated by drawing 10,000 random gene sets of same size from the dataset, subjecting the gene sets to principal component analysis and calculating the relative distance between the categories of the dimensionality reduced values.

## Results

### NRG-1/ERBB4 pathway controls STAT5b signaling in vivo in murine heart

To explore the role of STAT5b in ERBB4-mediated myocardial growth, mice expressing the soluble ectodomain (ECD) of ERBB4 in the heart as a result of infection with an adeno-associated virus of serotype 9 carrying the mouse ERBB4 ECD insert (AAV9-mERBB4ECD) were utilized. The ERBB4 ECD traps ERBB4 ligands, such as NRG-1, leading to downregulation of ERBB4 signaling in the cardiomyocytes.^28^

First, the effect of AAV9-mERBB4ECD treatment on cardiomyocyte size was assessed to learn whether reduction in ERBB4-sensitive signaling is associated with cardiomyocyte growth. The cardiomyocyte thickness was determined from heart sections (Figure 1A-B). As expected, the average cardiomyocyte thickness was lesser in the AAV9-mERBB4ECD-treated mice as compared to the controls (Figure 1A-B).

Activation of STAT5b by phosphorylation in the key tyrosine residue 699 leads to STAT5b dimerization and consequential accumulation into the nucleus at the target gene transcription sites. To explore the role of ErbB4 ligands in STAT5b activation in cardiomyocytes, the nuclear accumulation of STAT5b in the AAV9-mERBB4ECD and AAV9-control treated hearts was addressed by immunostaining (Figure 1C-D). The nuclear accumulation of STAT5b was found to be reduced in the AAV9-mERBB4ECD-treated hearts as compared to the controls, indicating a role for ERBB4 ligands in STAT5b activation in cardiomyocytes (Figure 1C-D).

The effect of the AAV9-mERBB4ECD-treatment on the transcription of known STAT5b target genes was further assessed with real time RT-PCR (Figure 1E). The treatment with AAV9-mERBB4ECD significantly decreased the levels of *Igf1* and *Myc* transcripts (Figure 1E), both of which are encoded by known STAT5b target genes that have been found to promote cardiac hypertrophy *in vivo* ^35–40^. In addition to *Igf1* and *Myc*, the transcription of a third STAT5b target gene, *Cdkn1a* ^41^, was similarly decreased.

### NRG-1/ERBB4 pathway controls STAT5b signaling in vitro in murine cardiomyocytes

To explore the STAT5b-dependency of NRG-1/ERBB4-mediated cardiomyocyte growth, cardiomyocytes from neonatal mice were isolated. As expected, the primary cardiomyocytes expressed ERBB4 JM-b more than the JM-a isoform (Supplementary Figure 1C), were positive for cardiomyocyte lineage specific markers myosin-heavy-chain and tropomyosin (Supplementary Figure 1D), and exhibited sporadic contractions in the culture plate. To assess the effect of STAT5b knock-down on NRG-1-induced cardiomyocyte hypertrophy, the isolated cardiomyocytes were infected with lentiviral particles containing plasmids encoding *Stat5b*-targeting shRNAs or control shRNA (Supplementary Figure 1E) and stimulated or not with NRG-1 for 2 days. The cardiomyocytes were stained with an antibody recognizing the heavy chain of a skeletal and cardiac myocyte specific myosin II, analyzed with immunofluorescence, and the cross-sectional area of the cardiomyocytes was quantified from confocal images taken in plane with the plasma membrane. The *Stat5b* knock-down significantly reduced the hypertrophic response induced by NRG-1 (Figure 1F-G), suggesting that NRG-1-stimulated hypertrophy is at least partially dependent on STAT5b.

To address the role of ERBB4 as a receptor for NRG-1 in mediating the hypertrophic response, *Erbb4* was knocked down in mouse primary cardiomyocytes by RNA interference (Supplementary Figure 1F-H). The treatment with *Erbb4*-targeting shRNAs significantly reduced the nuclear accumulation of STAT5b, as well as the cross-sectional area of cardiomyocytes (Figure 1H-I). To confirm that the transcription of STAT5b target genes in cardiomyocytes was STAT5b- and ERBB4-dependent, expression of *Igf1, Myc* and *Cdkn1a* was analyzed in cells subjected to *Stat5b* or *Erbb4* knock-down after stimulating the cells for 30 minutes with NRG-1. Real-time RT-PCR analyses demonstrated that both *Stat5b- and Erbb4-targeting* shRNAs significantly downregulated the expression of all three STAT5b target genes (Figure 1J). Taken together, these results demonstrate that the loss of NRG-1/ERBB4 signaling correlates with the loss of STAT5b signaling and reduced cardiomyocyte size in murine cardiomyocytes.

### Dynamin-2 controls subcellular ERBB4 localization and STAT5b activation

Mass spectrometry-derived ERBB4 interactome indicated that dynamin-2 (encoded by *DNM2* gene) was one of the prominent interaction partners of ERBB4 (Supplementary Figure 2A). Experimentation with proximity ligation assay (PLA) suggested that the heart-specific ERBB4 JM-b isoform was more prone to associate with dynamin-2 than the alternative isoform ERBB4 JM-a (Supplementary Figure 2B-C). The interaction between dynamin-2 and ERBB4 may have implications for heart biology, since the loss of *Dnm2* has been, similarly to the loss of *Nrg1* and *Erbb4*, associated with the development of heart failure ^42^. Previous research also indicates that the dynamin GTPases control vesicular endocytotic and ER-to-golgi trafficking. Of the three genes encoding dynamins, *DNM2* is most abundantly expressed in the heart (Supplementary Figure 2D).^42^

**Figure 2.**
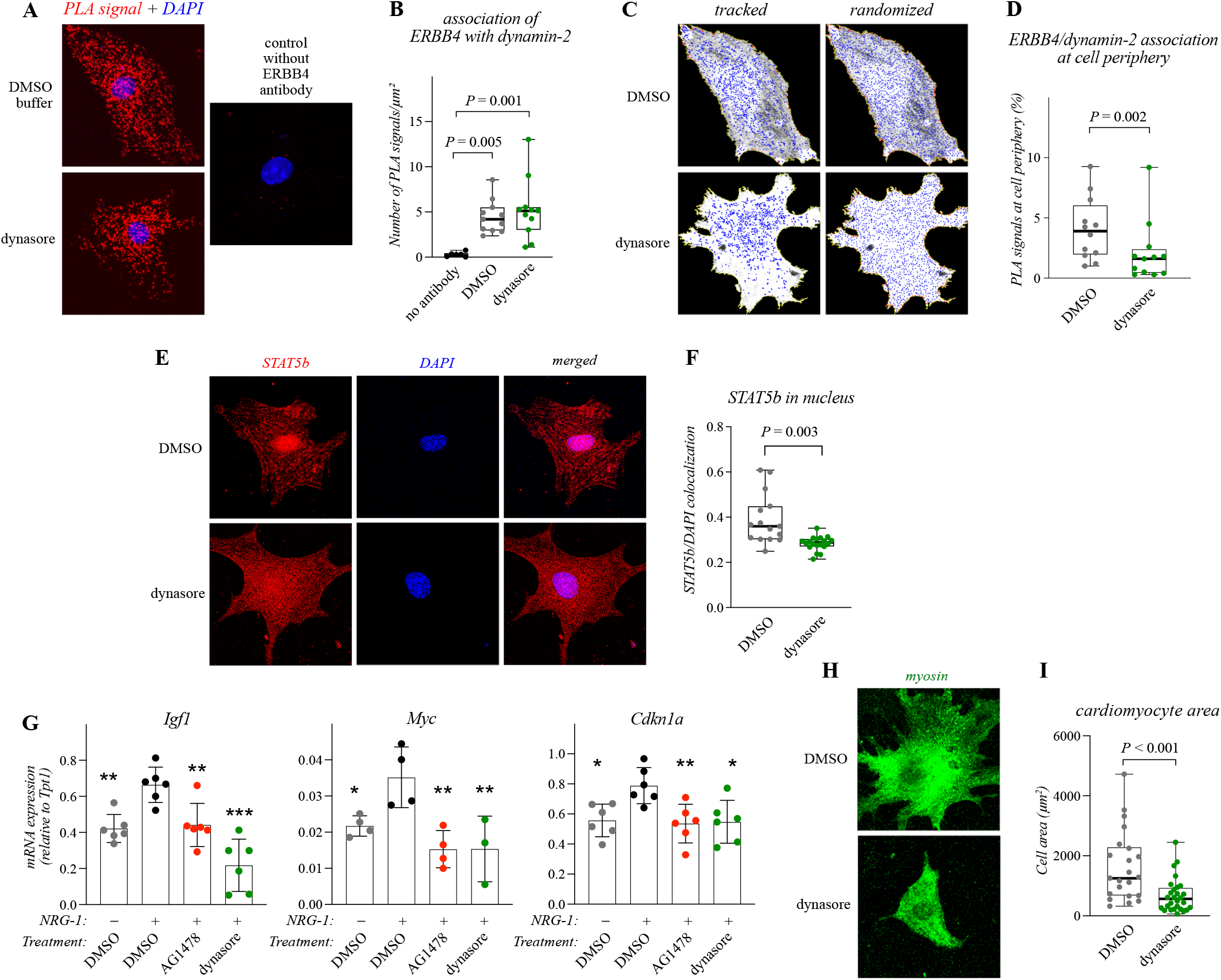
Dynamin-2 controls ERBB4 signaling and cardiomyocyte hypertrophy. **A-B**: Proximity ligation assay (PLA) of association of dynamin-2 with ERBB4 in primary mouse cardiomyocytes treated with either DMSO or the dynamin inhibitor dynasore for 5 hours. Panel A depicts a z projection of a stack of confocal images. PLA interactions are shown in red, nuclear stain DAPI in blue. Panel B depicts quantification of the data. One dot in the boxplot corresponds to the number of PLA signals in µm² (n = 6-11; combined from two replicate experiments). Non-parametric Kruskal-Wallis ANOVA and the Dunn’s multicomparison test with multiple test correction was used for statistics. **C-D:** Analysis of the subcellular localization of the association of dynamin-2 with ERBB4. Panel C depicts dynamin-2/ERBB4 PLA interactions in blue in representative cells (tracked) as well as when the PLA signals were artificially randomly distributed thoughout the area of the cell (randomized). Panel D depicts quantification of the PLA interactions within 1 µm distance from the edges of the cells (cell periphery). One dot in the boxplot corresponds to the fraction of PLA signals in the cell periphery in one cardiomyocyte (n = 12; combined from two replicate experiments). Non-parametric Mann-Whitney U-test was used. **E-F:** Confocal images (E) and quantification (F) of immunofluorescence staining of STAT5b and the nuclear stain DAPI in primary mouse cardiomyocytes. The cells were treated with either DMSO or dynasore for 5 hours and stimulated for 15 minutes with NRG-1. Panel F depicts quantification of colocalization of STAT5b- and DAPI-specific signals. One dot in the boxplots corresponds to one cell (n = 15-19; combined from two replicate experiments). Non-parametric Mann-Whitney U-test was used. **G:** Real-time RT-PCR analysis of expression of the indicated STAT5b target genes in primary mouse cardiomyocytes treated for 1-3 hours with the DMSO buffer, the ERBB kinase inhibitor AG1478, or dynasore, and stimulated or not for 30 minutes with NRG-1. One dot corresponds to the mean of technical repeats in one experiment and the whiskers the standard deviation (n = 3-6; replicate experiments). One-way ANOVA and the Dunnett’s multicomparison test with multiple test correction was used. *, *P* < 0.05; ** *P* < 0.01; ***, *P* < 0.001; against DMSO+ NRG-1 treatment. **H-I:** Confocal images (H) and quantification (I) of immunofluorescence staining of myosin heavy chain in primary mouse cardiomyocytes. The cells were treated with either DMSO or dynasore and stimulated with NRG-1 for two days. The myosin immunoreactivity was used to quantify cell area. One dot in the boxplot corresponds to one cell (n = 22-28; combined from three replicate experiments). Non-parametric Mann-Whitney U-test was used.

To test whether ERBB4 interacts with dynamin-2 in cardiomyocytes, the association of the two proteins was addressed with PLA in primary murine cardiomyocytes. In addition, to analyze whether the interaction was dependent on the dynamin-2 GTPase activity, the cardiomyocytes were cultured in the presence or absence of the dynamin GTPase inhibitor dynasore. ERBB4 and dynamin-2 indeed associated with each other in both control- and dynasore-treated cardiomyocytes (Figure 2A-B). Treatment with dynasore, however, significantly affected the subcellular localization of the associations between endogenously expressed ERBB4 and dynamin-2. Dynasore selectively reduced the associations at the cell surface and periphery of the cardiomyocytes (Figure 2C-D).

To further explore the effect of dynamin-inhibition on ERBB4 location, ERBB4 isoforms expressed in MCF-7 cells were analyzed for colocalization with the plasma membrane marker Na-K ATPase after treatment with dynasore or the control buffer. Dynasore significantly reduced the colocalization of both ERBB4 JM-a and JM-b isoforms with Na-K ATPase at the cell surface, indicating that the cell surface localization of ERBB4 was specifically dependent on the GTPase activity of dynamin-2 (Supplementary Figure 2E-F).

To explore whether the effect of dynamin-2 on ERBB4 localization was reflected in the downstream signaling via STAT5b, ERBB4 JM-b was overexpressed in mammary epithelial cells treated with dynasore or *DNM2*-targeting siRNAs. Both treatments abolished ERBB4 overexpression-promoted phosphorylation of the activating residue of STAT5b (Supplementary Figure 2G-H) and dynasore treatment additionally disrupted the co-precipitation of ERBB4 JM-b with STAT5b (Supplementary Figure 2I). Moreover, nuclear accumulation of STAT5b was reduced in cardiomyocytes treated with dynasore (Figure 2E-F), and both a chemical ERBB inhibitor AG1478 as well as dynasore significantly reduced the NRG-1-stimulated expression of STAT5b target genes (Figure 2G).

To control that the inhibition of the GTPase activity of dynamin-2 also affected the NRG-1-stimulated hypertrophic response, dynasore-treated cardiomyocytes were stimulated with NRG-1 for 2 days and analyzed for cross-sectional area. The area was significantly reduced by dynasore (Figure 2H-I). Taken together, the results indicate that dynamin-2 regulates subcellular ERBB4 localization and, consequently, NRG-1-stimulated STAT5b activation and cardiomyocyte growth.

### NRG-1 activates Stat5 in a zebrafish model of hyperplastic myocardial growth in vivo

In addition to hypertrophic growth, NRG-1/ErbB signaling has been reported to induce hyperplastic myocardial growth in zebrafish models ^8,43^. To explore the role of STAT5b also in the context of hyperplastic growth, NRG-1 or the control substance (BSA) was injected into the pericardial cavity of zebrafish embryos at 2 days post fertilization (dpf). The embryos were fixed at 4 dpf and stained with a myosin heavy chain antibody to visualize the myocardium. The cross-sectional ventricular area and ventricle thickness was measured from confocal images taken from anterior cross-sections of the ventricles. The NRG-1 injection induced hyperplastic cardiomyocyte growth as indicated by the increase in cross-sectional ventricular area but not in ventricular wall thickness (Figure 3A-B). This was further confirmed by assessing the ratio of the amount of nuclei in the ventricle to the ventricular area of the NRG-1-injected and BSA-injected embryos (Supplementary Figure 3). An increase in the number of nuclei in the myocardium of NRG-1-injected embryos was observed indicating proliferative growth.

**Figure 3.**
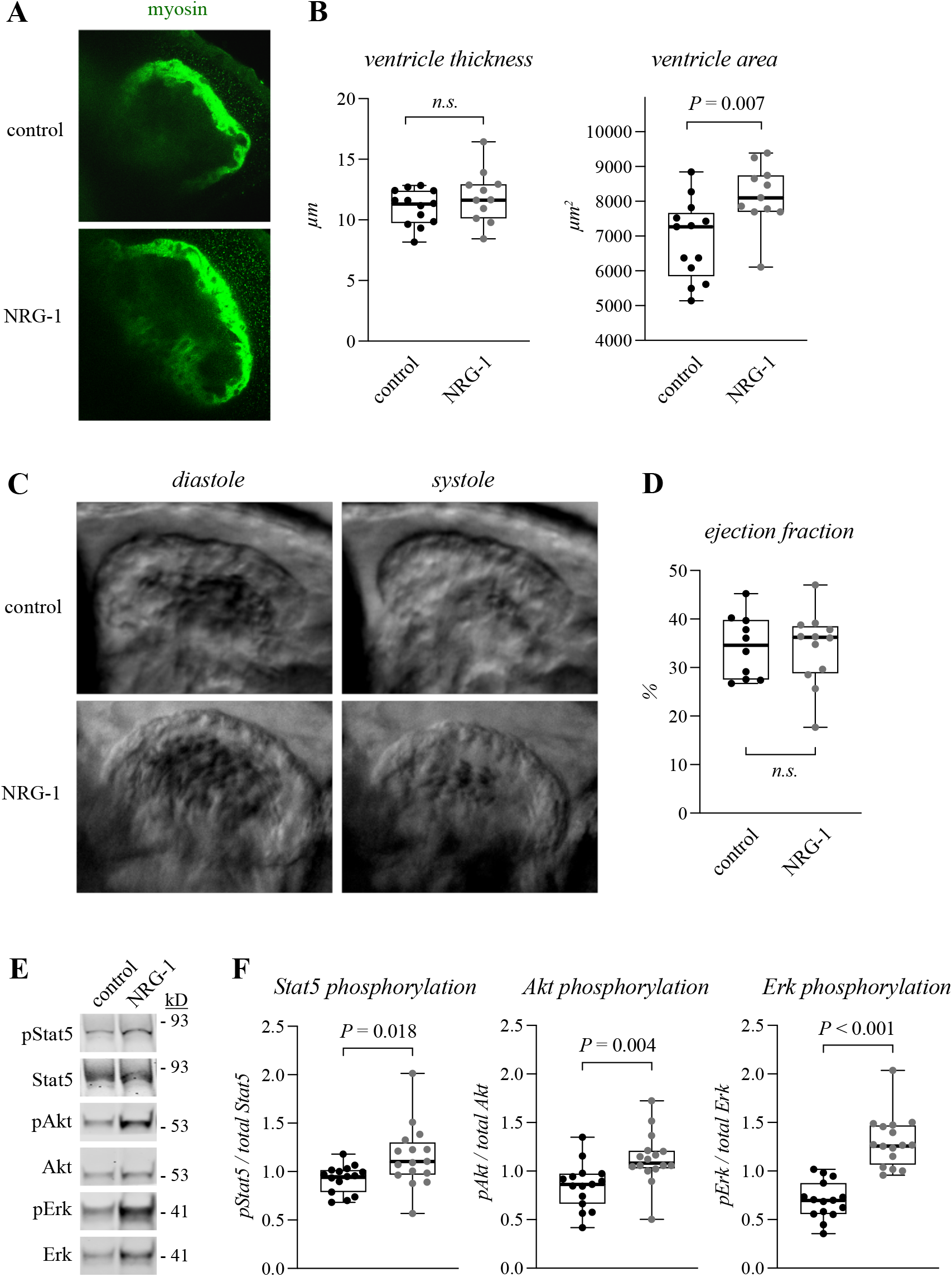
NRG-1 promotes myocardial growth and Stat5 activation in zebrafish embryos. **A-B:** Confocal images (A) and quantification (B) of immunofluorescence staining of myosin heavy chain in hearts of 4 dpf zebrafish embryos injected with either the buffer control or NRG-1 into the pericardial sac at 2 dpf. The myosin immunoreactivity was used to quantify both the average thickness of the ventricular wall as well as the cross-sectional area of the ventricles. One dot in the boxplots corresponds to one heart (n = 11-13; two replicate experiments). Unpaired two-tailed T-test was used for statistics. **C-D:** Still frame phase contrast images of in vivo imaging of the zebrafish embryo hearts (C) and quantification of ejection fractions (D). The zebrafish embryos were treated as in A. The ejection fractions were calculated from the videos. One dot in the boxplot corresponds to the ejection fraction of one heart (n = 10-12). Unpaired two-tailed T-test was used. **E-F:** Western analysis (E) and densitometric quantification (F) of Stat5, Akt and Erk phosphorylation in zebrafish embryos. The embryos were treated as in A. One dot in the boxplots corresponds to the relative densitometric value of one pooled sample of 3-4 zebrafish embryos (n = 15-16; combined from three replicate experiments). Unpaired two-tailed T-test with Welch’s correction (pStat5) or unpaired two-tailed T-test (pErk, pAkt) was used.

The effect of NRG-1 injection to the cardiac function of the zebrafish embryos was further assessed with live imaging (Figure 3C-D, Supplementary videos 1-2). The ventricles of the NRG-1-injected and BSA-injected zebrafish demonstrated similar ejection fractions indicating that the NRG-1 induced hyperplastic growth was physiological.

To assess the signaling pathways activated by the NRG-1 injections, western analyses were performed (Figure 3E and F). Unsurprisingly, the Pi3k/Akt and Erk pathways were activated with NRG-1. In addition, the Stat5 pathway was activated as indicated by the increased phosphorylation in the activating residue of the zebrafish Stat5 (Figure 3E and F). While the phospho-specific antibody does not differentiate between the zebrafish Stat5a and Stat5b, it is of note that both zebrafish Stat5a and Stat5b share more sequence similarity with human STAT5b than with human STAT5a. These data indicate that in a zebrafish model of NRG-1-induced hyperplastic myocardial growth, Stat5 is activated by the Nrg-1/Erbb4 pathway.

### Chemical inhibition of the Nrg-1/Erbb4 pathway results in reduced myocardial growth and Stat5 activation in vivo

To assess the effect of the Nrg-1/Erbb4 pathway inhibition on the hyperplastic myocardial growth and Stat5 activation *in vivo*, zebrafish embryos were treated at 2 dpf for 2 days with DMSO or the ERBB inhibitors AG1478, lapatinib or gefitinib. The function and the morphology of the ventricles was examined with live-imaging (Figure 4A). The ERBB inhibitor AG1478 has been reported to inhibit ERBB1, ERBB2 and ERBB4 ^44^, lapatinib ERBB1 and ERBB2 ^45^, and gefitinib ERBB1 ^46^. As expected, treatment with both inhibitors reported to inhibit the ERBB2/ERBB4 heterodimer in the myocardium significantly reduced the cross-sectional area of the ventricles, while treatment with the ERBB1 inhibitor gefitinib had no significant effect (Figure 4A-B). Treatment with AG1478 also reduced the thickness of the ventricular wall. The ejection fraction of all ERBB inhibitor-treated zebrafish, however, was retained although the AG1478- and lapatinib-treated ventricles displayed abnormal contractions (Supplementary videos 3-6).

**Figure 4.**
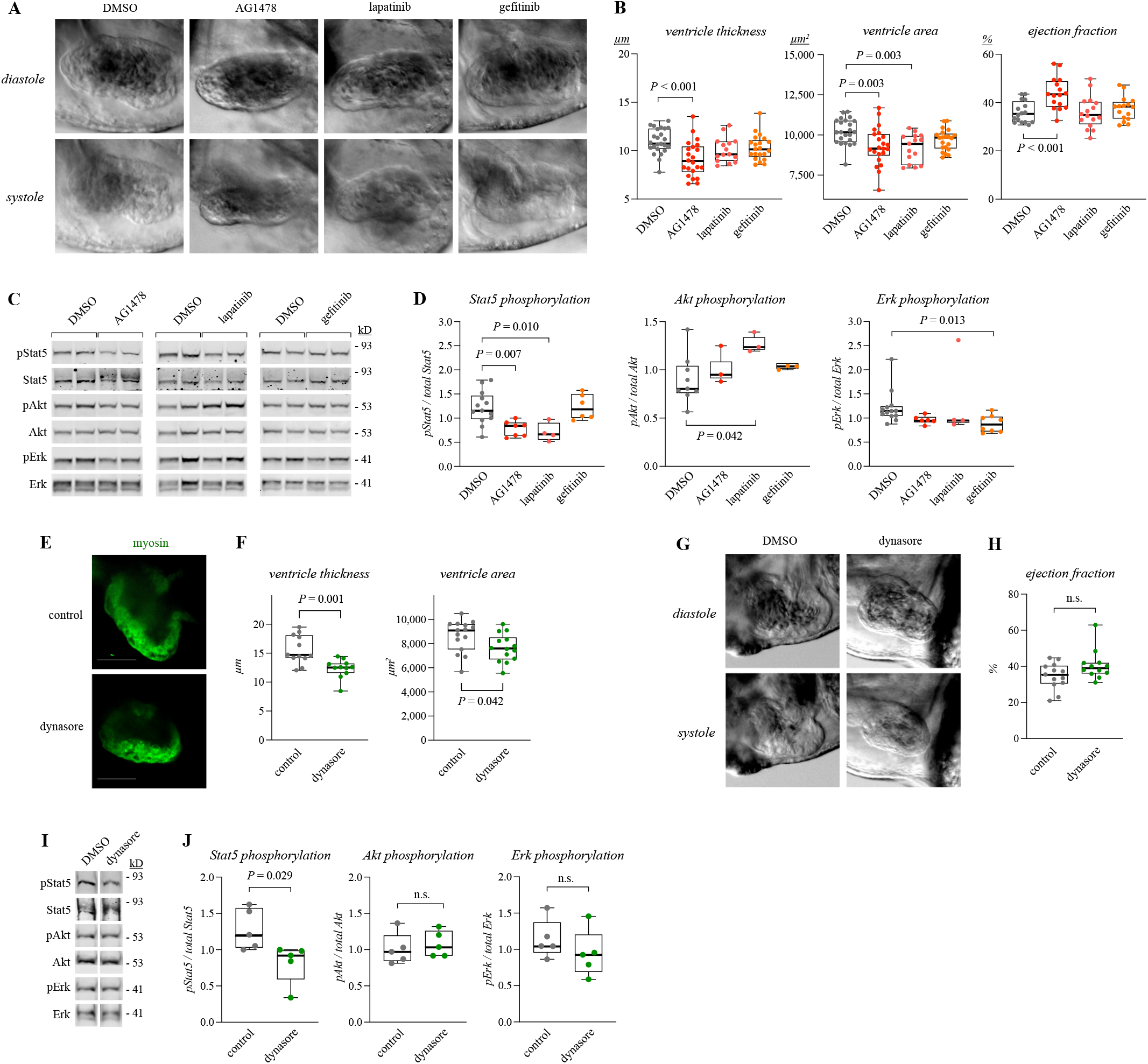
Erbb4 pathway regulates myocardial growth and Stat5 activation in zebrafish embryos. **A-B:** Still frame phase contrast images of in vivo imaging of the zebrafish embryo hearts (A) and quantification of heart dimensions and ejection fractions (B). The 4 dpf embryos were treated with the buffer control DMSO or the ERBB inhibitors AG1478, lapatinib or gefitinib for 2 days. The ventricular wall thickness, cross-sectional ventricular area, and ejection fraction was quantified from the videos. One dot in the boxplots corresponds to one heart (n = 16). One-way ANOVA and the Dunnett’s multicomparison test with multiple test correction was used for statistics. **C-D:** Western analysis (C) and densitometric quantification (D) of Stat5, Akt and Erk phosphorylation in zebrafish embryos. The embryos were treated as in A. One dot in the boxplots corresponds to the relative densitometric value of one pooled sample of 5 zebrafish embryos (n = 3-13; combined from three replicate experiments). One-way ANOVA and the Dunnett’s multicomparison test (pStat5, pAkt) or non-parametric Kruskal-Wallis with Dunn’s multicomparison test (pErk) was used. **E-F:** Confocal images (E) and quantification (F) of immunofluorescence staining of myosin heavy chain in hearts of 4 dpf zebrafish embryos treated with DMSO or dynasore for 2 days. The myosin immunoreactivity was used to quantify the average thickness of the ventricular wall and the cross-sectional area of the ventricles. One dot in the boxplots corresponds to one heart (n = 11-12). Unpaired two-tailed T-test was used. **G-H:** Still frame phase contrast images of in vivo imaging of the zebrafish embryo hearts (G) and quantification of ejection fractions (H). The zebrafish embryos were treated as in E. The ejection fractions were calculated from the videos. One dot in the boxplot corresponds to the ejection fraction of one heart (n = 12-13). Unpaired two-tailed T-test was used. **J:** Western analysis (I) and densitometric quantification (J) of Stat5, Akt and Erk phosphorylation in zebrafish embryos. The embryos were treated as in E. One dot in the boxplots corresponds to the relative densitometric value of one pooled sample of 5 zebrafish embryos (n = 5). Unpaired two-tailed T-test was used.

Consistent with the role of STAT5b downstream of ERBB4 activation in the mouse models (Figure 1C-D and H-I), treatment with the inhibitors targeting the ERBB2/ERBB4 heterodimer, but not the ERBB1 inhibitor, reduced Stat5 phosphorylation in zebrafish embryos (Figure 4C-D). Phosphorylation of Erk, however, was significantly reduced only by treatment with the ERBB1 inhibitor gefitinib. Signal for phospho-Akt, surprisingly, was induced by lapatinib and was not significantly affected by treatment with other ERBB inhibitors.

To examine the role of dynamin-2 in the hyperplastic growth and Stat5 activation, the embryos were treated at 2 dpf for 2 days with or without dynasore. The dynasore treatment resulted in both reduced ventricular area as well as ventricular wall thickness as compared to the control treatment (Figure 4E-F). The ejection fraction was, however, unaltered in the dynasore-treated ventricles indicating that the reduced myocardial growth did not lead to heart failure (Figure 4G-H, Supplementary videos 7-8).

Finally, the effect of dynasore on the activation of Stat5, Akt and Erk signaling was analyzed (Figure 4I-J). Stat5 was significantly less active in the dynasore-treated zebrafish embryos as compared to the control, while no effect was observed for either Akt or Erk signaling. Taken together the results indicate that the loss of myocardial growth due to inhibition of the Nrg-1/Erbb4/Dynamin-2 pathway correlates with the loss of Stat5, but not of Akt or Erk, activation.

### CRISPR/Cas9-mediated knock-down of the zebrafish stat5b gene results in reduced myocardial growth and heart failure

To explore whether knock-down of the *stat5b* gene was sufficient to reduce myocardial growth, zebrafish embryos were injected with the recombinant CRISPR/Cas9 enzyme and control- or *stat5b*-targeting guide RNAs (gRNA) at one-cell stage. The efficacy of the *stat5b* knock-down was confirmed with genomic sequencing, real-time RT-PCR (Supplementary Figure 4) and with immunofluorescence staining of whole-mount embryos (Figure 5A-B). The cross-sectional area of the ventricles of *stat5b* gRNA-injected 4 dpf embryos was significantly smaller when compared to the control gRNA-injected embryos (Figure 5A-B), indicating reduced myocardial growth.

**Figure 5.**
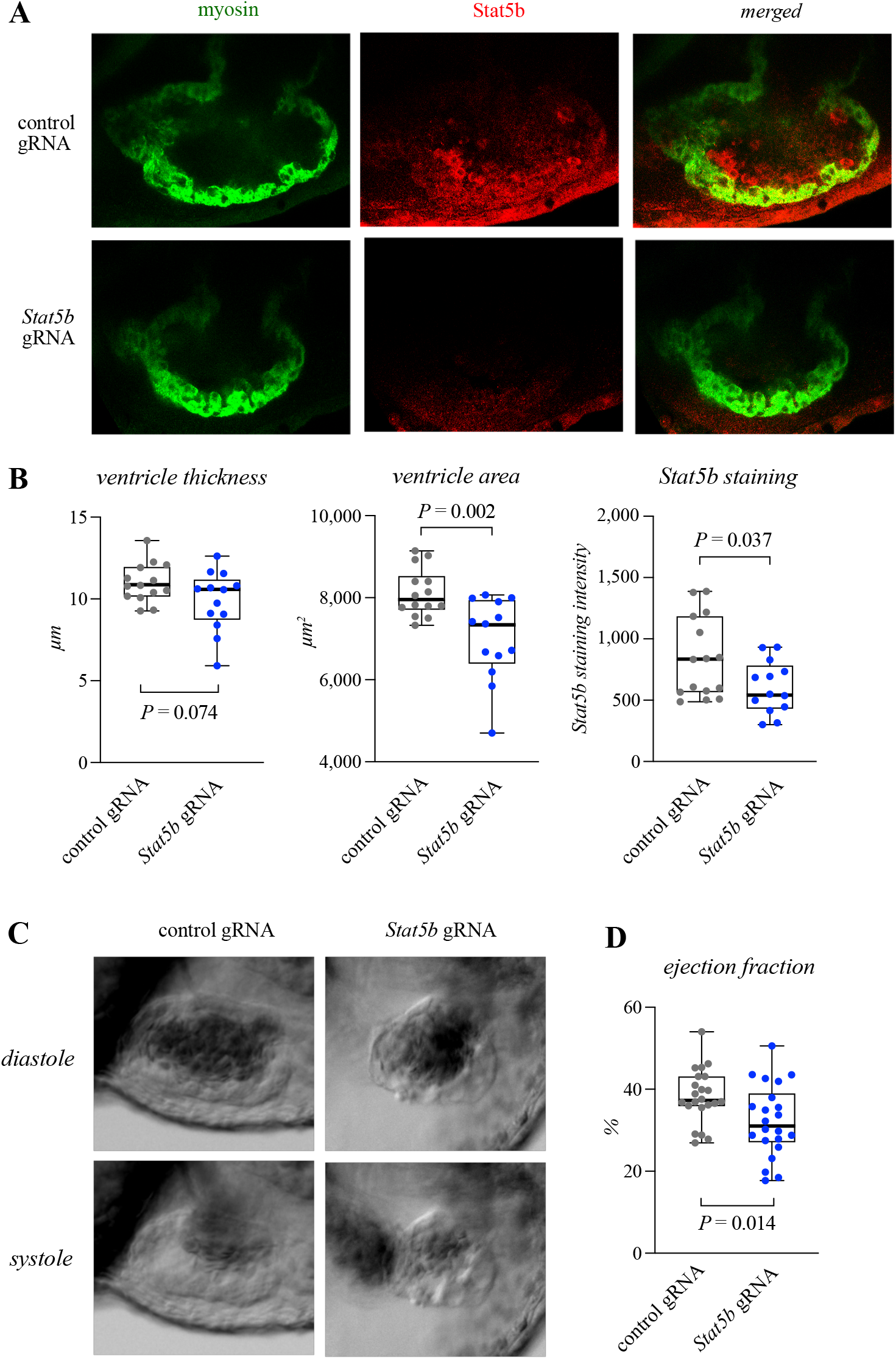
Stat5b is necessary for cardiac growth and function in zebrafish embryos. **A-B:** Confocal images (A) and quantification (B) of immunofluorescence staining of myosin heavy chain and Stat5b in hearts of 4 dpf zebrafish embryos injected with CRISPR/Cas9 and either control or STAT5b-targeting gRNA at one-cell stage. The myosin immunoreactivity was used to quantify the average thickness of the ventricular wall and the cross-sectional area of the ventricles. Stat5b staining intensity was also quantified. One dot in the boxplots corresponds to one heart (n = 13-14; combined from two replicate experiments). Unpaired two-tailed T-test was used for statistics. **C-D:** Still frame phase contrast images of in vivo imaging of the zebrafish embryo hearts (C) and quantification of the ejection fraction (D). The 4 dpf embryos were treated as in A. Ejection fraction was quantified from the videos. One dot in the boxplots corresponds to ejection fraction of one heart (n = 22; two replicate experiments). Unpaired two-tailed T-test was used.

The ejection fraction of the ventricles of *stat5b* gRNA-injected zebrafish embryos was also significantly reduced compared to the control gRNA-injected embryos (Figure 5C-D; Supplementary videos 9-10) suggesting heart failure. These results indicate that the downregulation of Stat5b alone is sufficient to disrupt the normal growth and function of the myocardium.

### The NRG-1/ERBB4/STAT5b signaling pathway is dysregulated in the myocardium of patients with pathological cardiac hypertrophy

Finally, to explore the role of the NRG-1/ERBB4/STAT5b signaling pathway in the context of pathological hypertrophic myocardial growth, the expression signature of the NRG-1/ERBB4/STAT5b signaling pathway and activation of STAT5b were investigated in clinical samples representing different types of pathological cardiac hypertrophy or normal heart tissue. First, pre-existing transcriptome datasets were subjected to principal component analysis to observe whether control subjects and patients suffering from hypertrophic cardiomyopathy could be segregated based on the expression levels of the genes in the NRG-1/ERBB4/STAT5b signaling pathway (Figure 6A; Supplementary Figure 5A). The details of the datasets can be accessed through the gene expression omnibus database ^33,34^ with the GSE prefix identifiers indicated in the figure legends of Figure 6 and in the Supplementary Figure 5. The dimensionality reduction of the expression level of the NRG-1/ERBB4/STAT5b signaling pathway genes indeed was able to significantly cluster cases with pathological cardiac hypertrophy and those without into separate groups in three independent datasets. The successful segregation indicates that the NRG-1/ERBB4/STAT5b signaling pathway genes are consistently differentially regulated in hypertrophic vs. normal cardiac tissue. The transcripts of *MYC, IGF1* and *NRG1* genes were most differentially regulated in the NRG-1/ERBB4/STAT5b signaling pathway, as indicated by the principal component weights (Supplementary Figure 5B).

**Figure 6.**
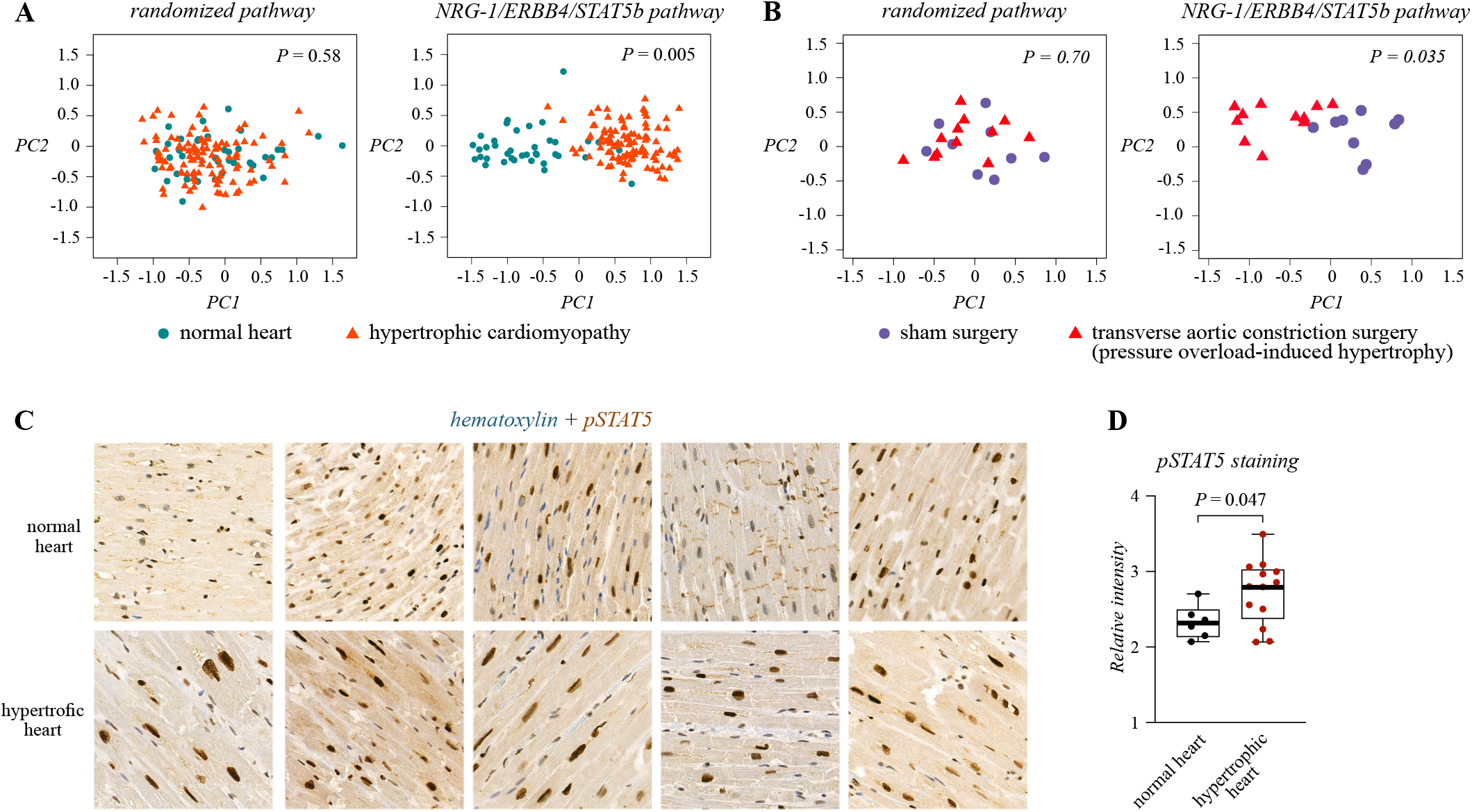
NRG-1/ERBB4/STAT5b signaling pathway is perturbed in pathological cardiac hypertrophy. **A:** Principal component and cluster analysis of transcripts of the NRG-1/ERBB4/STAT5b pathway (including the genes: *NRG1, ERBB4, STAT5B, IGF1, MYC* and *DNM2*) with clinical samples representing normal myocardium or hypertrophic cardiomyopathy. The dataset was acquired from the GEO database (identifier GSE36961). One symbol in the plots corresponds to one subject (n = 39-106). Statistical significance of clusterization was calculated by estimating a probability distribution for the relative distance within a cluster and between clusters and by drawing the cumulative probability from the resulting empirical cumulative probability function. **B:** Principal component and cluster analysis of transcripts of the NRG-1/ERBB4/STAT5b pathway with samples representing myocardia of mice subjected to sham or transverse aortic constriction surgery. The dataset was acquired from the GEO database (identifier GSE5500). One symbol in the plots corresponds to one mouse (n = 9-12). **C-D:** Immunohistochemical analysis (C) and quantification (D) of STAT5 activation in sections representing normal myocardia or pathological cardiac hypertrophy (n = 3, aortic stenosis; n = 2, idiopathic cardiomyopathy). Phospho-STAT5b staining intensity was quantified. One dot in the boxplot corresponds to one subject (n = 6-13). Unpaired two-tailed T-test was used for statistics.

The role of NRG-1/ERBB4/STAT5b signaling pathway in pressure overload-induced pathological cardiac hypertrophy was also investigated from datasets representing cardiac transcriptomes of mice that had undergone transverse aortic constriction surgery (TAC) or sham surgery. The dimensionality reduction of the expression level of the NRG-1/ERBB4/STAT5b signaling pathway genes led to a statistically significant clustering of the TAC- and sham-treated mice into separate groups in three independent datasets (Figure 6B; Supplementary Figure 5C). The successful segregation indicates that NRG-1/ERBB4/STAT5b signaling pathway genes are also differentially regulated in pressure overload-induced cardiac hypertrophy. The principal component weights suggested that *igf1* and *dnm2* were most differentially regulated by the TAC vs. sham surgery (Supplementary Figure 5D).

The activation of STAT5b in control subjects and patients suffering from pathological cardiac hypertrophy due to aortic stenosis, alcoholic cardiomyopathy or idiopathic cardiomyopathy was investigated with immunohistochemistry. Formalin-fixed paraffin-embedded (FFPE) sections from cardiac left ventricle were acquired from the Auria Biobank (University Hospital of Turku, Finland). Both heart weight and cardiomyocyte size were greater in the samples representing pathological cardiac hypertrophy (Supplementary Figure 5E). The heart sections were stained with an antibody that recognizes the phosphorylated state of the activating residue of STAT5b and the immunosignal was quantified from cardiomyocytes in representative randomly selected regions. The specificity of the immunosignal was controlled in FFPE samples treated with insulin or control or STAT5 targeting siRNAs (Supplementary Figure 6). The signal of the phosphorylated STAT5 in cardiomyocyte nuclei was significantly greater in the cases representing pathological cardiac hypertrophy as compared to the controls (Figure 6C-D; only five cases from both cohorts visualized) indicating that STAT5b activation is up-regulated in pathologically hypertrophic myocardium in human. Taken together, these results suggest that the NRG-1/ERBB4/STAT5b pathway is dysregulated in mammalian pathological cardiac hypertrophy.

## Discussion

A recent review identified the effect of NRG-1 on cardiomyocyte growth as one of the translationally relevant but poorly understood aspects of cardiovascular research, and called for further research on the underlying molecular mechanisms^47^. Here we focused on the signaling pathway of NRG-1 in primary cardiomyocytes as well as in the heart tissue of mouse and zebrafish to elucidate the molecular mechanisms of NRG-1-mediated cardiomyocyte growth. The NRG-1 receptor ERBB4, the ERBB4-activated signal transducer STAT5b, and dynamin-2, that was discovered to control the subcellular localization of ErbB4, were identified as key effectors controlling the expression of two transcription factors known as essential regulators of cardiac growth, IGF1 and MYC.

In our models, STAT5b mediated both hypertrophic and hyperplastic cardiomyocyte growth. In primary mouse cardiomyocytes and in the AAV9-mERBB4ECD-treated mice, NRG-1 induced a hypertrophic response while in zebrafish embryos the response was hyperplastic. These observations are consistent with previous reports of NRG-1 responses in murine and zebrafish *in vivo* models and in primary cardiomyocytes^1,2,43,48^. In accordance, the target genes of STAT5b *IGF1* and *MYC* have been associated with hypertrophic^39^ and hyperplastic^49^ growth, respectively, implying that the STAT5b mediated hypertrophic growth may be mediated by the expression of MYC and the hyperplastic growth by the expression of IGF1.

The ERK pathway has been established as one of the main pathways regulating hypertrophic cardiomyocyte growth downstream of several stimuli^50^. The NRG-1/ERBB pathway has also been shown to promote ERK1/2 phosphorylation in cardiomyocytes^2,15,17^. A recent report, however, suggests that the activation of ERK signaling alone is not sufficient to confer NRG-s1-mediated cardiomyocyte hypertrophy^22^. This is consistent with several reports where treatment with an anti-ERBB2 antibody or *ERBB2*-targeting siRNAs has been unable to reduce the phosphorylation of ERK1/2 in cardiomyocytes^15,22,51^. Treatment with *ERBB4* shRNA has drastically reduced hypertrophic growth but similarly has only led to a slight reduction of ERK1/2 phosphorylation in cardiomyocytes^22^.

While the conditional double knock-out of *Erk1/Erk2* in mice leads to dilated cardiomyopathy resembling the loss of the genes along the NRG-1/ERBB2/ERBB4 pathway ^52^, the histology of the dilated ventricles is markedly different between the conditional double *Erk1/Erk2* knock-out model and the conditional *Nrg1*/*Erbb2*/*Erbb4* pathway knock-out models. The cardiomyocytes in the double *Erk1/Erk2* knock-out model are elongated and show a loss of concentric hypertrophic response^52^, while the cardiomyocytes in the *Erbb2/Erbb4* knock-out models are hypertrophic, although the gross ventricular phenotype is dilated in both models ^10,11^. The asymmetry in the cardiomyocyte morphology of the *Erk1/Erk2* vs *Erbb2/Erbb4* conditional knock-out model hearts is consistent with our observations here of NRG-1/ERBB4 additionally activating STAT5b in murine *in vitro* and *in vivo* models of hypertrophic growth and the reduction of NRG-1/ERBB4-mediated hypertrophic cardiomyocyte growth by STAT5b knock-down.

The NRG-1-mediated hyperplastic growth has been attributed to PI3K/AKT signaling based on an experiment, in which the expression of PTEN, the negative regulator of PI3K signaling, reduced DNA synthesis ^1^. A more direct connection between hyperplastic cardiomyocyte growth and NRG-1-induced AKT signaling remains to be shown. In the results presented here, the activation of STAT5b was consistently modulated by intervention of the NRG-1/ERBB4 signaling pathway, while the phosphorylation of ERK and AKT was significantly modulated by NRG-1 injections but not by chemical inhibition of the NRG-1/ERBB4 pathway in zebrafish models of hyperplastic myocardium growth.

Similar to NRG-1 and ERBB4 ^47^, STAT5b has previously been proposed to have role in the molecular pathogenesis of heart failure. Indeed, a polymorphism in the exon 1 of the *STAT5B* gene has been associated with the risk for dilated cardiomyopathy^53^. Here we identified differential regulation of the NRG-1/ERBB4/DNM2/STAT5b signaling in clinical samples of pathological cardiac hypertrophy independent of the etiology. This is consistent with previous observations that chemical inhibition of STAT5 reduces pathological cardiac hypertrophy in *in vivo* mouse models^23,54^. Pathological cardiac hypertrophy is a compensatory response that can precede heart failure and ERBB4 expression has been reported to be downregulated in cases in which cardiac hypertrophy has advanced to heart failure ^55,56^. In our zebrafish model, the CRISPR/Cas9-mediated knock-down of *stat5b* also led to reduced ejection fraction indicative of heart failure. Further experimentation is, however, needed to understand the role of STAT5b in pathological cardiac hypertrophy and its progression to heart failure.

STAT5b has been suggested to mediate the cardioprotective effect of remote ischemic preconditioning in a murine model of ischemic cardiac injury as well as in clinical samples ^57–59^. The results presented here further support the therapeutic potential of STAT5b by introducing STAT5b as one of the downstream effectors of NRG-1 in the heart. NRG-1 signaling has been found to attenuate cardiac injury in several animal models including e.g. ischemia, doxorubicin, metabolic stress and pacing induced cardiac injury models ^60,61^. The cardioprotective effect of NRG-1 has been attributed to increased PI3K/AKT signaling ^60,61^. Interestingly in a murine model of ischemic cardiac injury remote ischemic preconditioning increased the activation of PI3K/AKT signaling only in the presence of STAT5b, suggesting that the cardioprotective PI3K/AKT signaling is activated downstream of STAT5b ^57^. Consistently, chemical inhibition of STAT5 has been shown to downregulate PI3K/AKT signaling in a mouse model of pressure overload induced cardiac hypertrophy^54^. It remains to be formally addressed whether the cardioprotective effect of NRG-1 is dependent on STAT5b and whether e.g. targeting the deactivating phosphatases of STAT5b^62–64^ or repressors of STAT5b induced transcription^65–67^ in the myocardium could serve as an approach with therapeutic potential in heart failure. In accordance with clinical applicability, administration of NRG-1 and the protein product of the STAT5b target gene *IGF1* has demonstrated success in attenuating dilated cardiomyopathy in clinical trials^68–70^.

In addition to STAT5b, the membrane-remodeling GTPase dynamin-2 was identified as a regulator of NRG-1/ERBB4 signaling promoting cardiomyocyte growth. Inhibition of dynamin GTPase activity reduced ERBB4 at the cell surface as well as STAT5b activation and NRG-1-induced cardiomyocyte and myocardial growth in zebrafish embryos. Expression of dynamin-2 has previously been reported to be decreased in heart failure in isopretenol-induced rat heart failure model and in clinical samples ^42^. It has also been demonstrated that decreased dynamin-2 expression leads to cardiomyocyte apoptosis by stimulating Ca^2+^ influx from the sarcolemma ^42^. Interestingly, several reports also indicate a role for NRG-1/ERBB4 signaling in regulating cardiomyocyte apoptosis ^71–73^, suggesting that cardiomyocyte apoptosis induced by dynamin-2 deficiency ^42^ could be mediated by the loss of NRG-1/ERBB4 signaling.

## Funding

This work was supported by the Academy of Finland, Cancer Foundation Finland, the Sigrid Juselius Foundation, the Turku University Central Hospital, Orion Research Foundation, K. Albin Johanssons Foundation, the Cancer Society of Southwestern Finland, the Maud Kuistila memorial Foundation, Emil Aaltonen Foundation, the Finnish Foundation for Cardiovascular Research, Paavo Nurmi Foundation, Finnish Cultural Foundation and the Varsinais-Suomi Regional Fund of the Finnish Cultural Foundation.

## Acknowledgements

We thank Maria Tuominen, Mika Savisalo, Nea Konttinen, Merja Lakkisto, Minna Santanen and Sinikka Kollanus for skillful technical assistance, and Riku Kiviranta and his laboratory for providing research material. Turku Doctoral Program of Molecular Medicine is acknowledged for excellent doctoral training and funding. The Auria biobank is acknowledged for providing patient samples. We acknowledge the Cell Imaging Core, Proteomics Core, Zebrafish Core and Finnish Functional Genomics Centre (Turku Bioscience Centre, University of Turku and Åbo Akademi University) all supported by Biocenter Finland and Histology Core Facility of the Institute of Biomedicine, University of Turku for their excellent service.

## Conflict of interests

The authors declare that they have no conflict of interests.

## Author contributions

KV and KE designed the study. KV, AJ and IP performed the experimentation with the help of JAM. KV performed statistical and data analysis. KAH, RK and KA provided the samples from AAV treated mice. PT was crucial in the acquirement, selection and staining of the patient samples. JH provided the instructions for ejection fraction calculation. KV and KE wrote the manuscript with the help of AJ and IP.

## References

1. Bersell K, Arab S, Haring B, Kühn B. Neuregulin1/ErbB4 Signaling Induces Cardiomyocyte Proliferation and Repair of Heart Injury. Cell 2009;138:257–270.

2. Baliga RR, Pimental DR, Zhao YY, Simmons WW, Marchionni MA, Sawyer DB, Kelly RA. NRG-1-induced cardiomyocyte hypertrophy. Role of PI-3-kinase, p70(S6K), and MEK-MAPK-RSK. Am J Physiol - Hear Circ Physiol 1999;277.

3. Lee KF, Simon H, Chen H, Bates B, Hung MC, Hauser C. Requirement for neuregulin receptor erbB2 in neural and cardiac development. Nature 1995;378:394–398.

4. Gassmann M, Casagranda F, Orloli D, Simon H, Lai C, Kleint R, Lemke G. Aberrant neural and cardiac development in mice lacking the ErbB4 neuregulin receptor. Nat 1995 3786555 1995;378:390–394.

5. Meyer D, Birchmeier C. Multiple essential functions of neuregulin in development. Nat 1995 3786555 1995;378:386–390.

6. Plowman GD, Green JM, Culouscou JM, Carlton GW, Rothwell VM, Buckley S. Heregulin induces tyrosine phosphorylation of HER4/p180erbB4. Nature 1993;366:473–475.

7. Gassmann M, Casagranda F, Orioli D, Simon H, Lai C, Klein R, Lemke G. Aberrant neural and cardiac development in mice lacking the ErbB4 neuregulin receptor. Nature 1995;378:390–394.

8. Liu J, Bressan M, Hassel D, Huisken J, Staudt D, Kikuchi K, Poss KD, Mikawa T, Stainier DYR. A dual role for ErbB2 signaling in cardiac trabeculation. Development 2010;137:3867–3875.

9. Crone SA, Zhao YY, Fan L, Gu Y, Minamisawa S, Liu Y, Peterson KL, Chen J, Kahn R, Condorelli G, Ross J, Chien KR, Lee KF. ErbB2 is essential in the prevention of dilated cardiomyopathy. Nat Med 2002;8:459–465.

10. Özcelik C, Erdmann B, Pilz B, Wettschureck N, Britsch S, Hübner N, Chien KR, Birchmeier C, Garratt AN. Conditional mutation of the ErbB2 (HER2) receptor in cardiomyocytes leads to dilated cardiomyopathy. Proc Natl Acad Sci U S A 2002;99:8880–8885.

11. García-Rivello H, Taranda J, Said M, Cabeza-Meckert P, Vila-Petroff M, Scaglione J, Ghio S, Chen J, Lai C, Laguens RP, Lloyd KC, Hertig CM. Dilated cardiomyopathy in Erb-b4-deficient ventricular muscle. Am J Physiol - Hear Circ Physiol 2005;289.

12. Gao R, Zhang J, Cheng L, Wu X, Dong W, Yang X, Li T, Liu X, Xu Y, Li X, Zhou M. A Phase II, randomized, double-blind, multicenter, based on standard therapy, placebo-controlled study of the efficacy and safety of recombinant human neuregulin-1 in patients with chronic heart failure. J Am Coll Cardiol 2010;55:1907–1914.

13. Jabbour A, Hayward CS, Keogh AM, Kotlyar E, McCrohon JA, England JF, Amor R, Liu X, Li XY, Zhou MD, Graham RM, MacDonald PS. Parenteral administration of recombinant human neuregulin-1 to patients with stable chronic heart failure produces favourable acute and chronic haemodynamic responses. Eur J Heart Fail 2011;13:83–92.

14. Zensun Sci. & Tech. Co. L. Survival Study of the Recombinant Human Neuregulin-1β in Subjects With Chronic Heart Failure. https://clinicaltrials.gov/ct2/show/NCT03388593https://clinicaltrials.gov/ct2/show/NCT03388593 (8 March 2022)

15. Kuramochi Y, Guo X, Sawyer DB. Neuregulin activates erbB2-dependent src/FAK signaling and cytoskeletal remodeling in isolated adult rat cardiac myocytes. J Mol Cell Cardiol 2006;41:228.

16. Pentassuglia L, Heim P, Lebboukh S, Morandi C, Xu L, Brink M. Neuregulin-1␤ promotes glucose uptake via PI3K/Akt in neonatal rat cardiomyocytes. Am J Physiol Endocrinol Metab 2016;310:E782–E794.

17. Fukazawa R, Miller TA, Kuramochi Y, Frantz S, Kim Y-D, Marchionni MA, Kelly RA, Sawyer DB. Neuregulin-1 protects ventricular myocytes from anthracycline-induced apoptosis via erbB4-dependent activation of PI3-kinase/Akt. J Mol Cell Cardiol 2003;35:1473–1479.

18. Jie B, Zhang X, Wu X, Xin Y, Liu Y, Guo Y. Neuregulin-1 suppresses cardiomyocyte apoptosis by activating PI3K/Akt and inhibiting mitochondrial permeability transition pore. Mol Cell Biochem 2012;370:35–43.

19. Cai M-X, Shi X-C, Chen T, Tan Z-N, Lin Q-Q, Du S-J, Tian Z-J. Exercise training activates neuregulin 1/ErbB signaling and promotes cardiac repair in a rat myocardial infarction model. 2016.

20. Vaparanta K, Jokilammi A, Tamirat M, Merilahti JAM, Salokas K, Varjosalo M, Ivaska J, Johnson MS, Elenius K. An extracellular receptor tyrosine kinase motif orchestrating intracellular STAT activation. submitted 2022.

21. Elenius K, Corfas G, Paul S, Choi CJ, Rio C, Plowman GD, Klagsbrun M. A Novel Juxtamembrane Domain Isoform of HER4/ErbB4: ISOFORM-SPECIFIC TISSUE DISTRIBUTION AND DIFFERENTIAL PROCESSING IN RESPONSE TO PHORBOL ESTER. J Biol Chem 1997;272:26761–26768.

22. Wang Z, Chan HW, Gambarotta G, Smith NJ, Purdue BW, Pennisi DJ, Porrello ER, O’Brien SL, Reichelt ME, Thomas WG, Paravicini TM. Stimulation of the four isoforms of receptor tyrosine kinase ErbB4, but not ErbB1, confers cardiomyocyte hypertrophy. J Cell Physiol 2021;236:8160–8170.

23. Jin G, Wang L, Ma J. Inhibiting STAT5 significantly attenuated Ang II-induced cardiac dysfunction and inflammation. Eur J Pharmacol 2022;915:174689.

24. Maatta JA, Sundvall M, Junttila TT, Peri L, Laine VJO, Isola J, Egeblad M, Elenius K. Proteolytic Cleavage and Phosphorylation of a Tumor-associated ErbB4 Isoform Promote Ligand-independent Survival and Cancer Cell Growth. Mol Biol Cell 2005;17:67–79.

25. Louch WE, Sheehan KA, Wolska BM. Methods in cardiomyocyte isolation, culture, and gene transfer. J Mol Cell Cardiol 2011;51:288–298.

26. White RM, Sessa A, Burke C, Bowman T, LeBlanc J, Ceol C, Bourque C, Dovey M, Goessling W, Burns CE, Zon LI. Transparent adult zebrafish as a tool for in vivo transplantation analysis. Cell Stem Cell 2008;2:183–189.

27. Nüsslein-Volhard C (Christiane), Dahm R. Zebrafish : a practical approach. 2002:303.

28. Kivelä R, Hemanthakumar KA, Vaparanta K, Robciuc M, Izumiya Y, Kidoya H, Takakura N, Peng X, Sawyer DB, Elenius K, Walsh K, Alitalo K. Endothelial Cells Regulate Physiological Cardiomyocyte Growth via VEGFR2-Mediated Paracrine Signaling. Circulation 2019;139:2570–2584.

29. Dull T, Zufferey R, Kelly M, Mandel RJ, Nguyen M, Trono D, Naldini L. A third-generation lentivirus vector with a conditional packaging system. J Virol 1998;72:8463–8471.

30. Hoshijima K, Jurynec MJ, Klatt Shaw D, Jacobi AM, Behlke MA, Grunwald DJ. Highly Efficient CRISPR-Cas9-Based Methods for Generating Deletion Mutations and F0 Embryos that Lack Gene Function in Zebrafish. Dev Cell 2019;51:645-657.e4.

31. Lund FW, Jensen ML V., Christensen T, Nielsen GK, Heegaard CW, Wüstner D. SpatTrack: An Imaging Toolbox for Analysis of Vesicle Motility and Distribution in Living Cells. Traffic 2014;15:1406–1429.

32. Villalta JI, Galli S, Iacaruso MF, Antico Arciuch VG, Poderoso JJ, Jares-Erijman EA, Pietrasanta LI. New Algorithm to Determine True Colocalization in Combination with Image Restoration and Time-Lapse Confocal Microscopy to Map Kinases in Mitochondria. Polymenis M, ed. PLoS One 2011;6:e19031.

33. Edgar R, Domrachev M, Lash AE. Gene Expression Omnibus: NCBI gene expression and hybridization array data repository. Nucleic Acids Res 2002;30:207–210.

34. Barrett T, Wilhite SE, Ledoux P, Evangelista C, Kim IF, Tomashevsky M, Marshall KA, Phillippy KH, Sherman PM, Holko M, Yefanov A, Lee H, Zhang N, Robertson CL, Serova N, Davis S, Soboleva A. NCBI GEO: archive for functional genomics data sets--update. Nucleic Acids Res 2013;41.

35. Basham B, Sathe M, Grein J, McClanahan T, D’Andrea A, Lees E, Rascle A. In vivo identification of novel STAT5 target genes. Nucleic Acids Res 2008;36:3802.

36. Lord JD, McIntosh BC, Greenberg PD, Nelson BH. The IL-2 Receptor Promotes Lymphocyte Proliferation and Induction of the c-myc, bcl-2, and bcl-x Genes Through the trans-Activation Domain of Stat5. J Immunol 2000;164:2533–2541.

37. Nosaka T, Kawashima T, Misawa K, Ikuta K, Mui ALF, Kitamura T. STAT5 as a molecular regulator of proliferation, differentiation and apoptosis in hematopoietic cells. EMBO J 1999;18:4754.

38. Rotwein P. Mapping the Growth Hormone – Stat5b – IGF-I Transcriptional Circuit. Trends Endocrinol Metab 2012;23:186.

39. Zhong W, Mao S, Tobis S, Angelis E, Jordan MC, Roos KP, Fishbein MC, Alborán IM De, MacLellan WR. Hypertrophic growth in cardiac myocytes is mediated by Myc through a Cyclin D2-dependent pathway. EMBO J 2006;25:3869–3879.

40. Delaughter MC, Taffet GE, Fiorotto ML, Entman ML, Schwartz RJ. Local insulin-like growth factor I expression induces physiologic, then pathologic, cardiac hypertrophy in transgenic mice. FASEB J 1999;13:1923–1929.

41. Yu JH, Zhu BM, Wickre M, Riedlinger G, Chen W, Hosui A, Robinson GW, Hennighausen L. The transcription factors STAT5A and STAT5B negatively regulate cell proliferation through the activation of Cdkn2b and Cdkn1a expression. Hepatology 2010;52:1808.

42. Li J, Zhang D-S, Ye J-C, Li C-M, Qi M, Liang D-D, Xu X-R, Xu L, Liu Y, Zhang H, Zhang Y-Y, Deng F-F, Feng J, Shi D, Chen J-J, Li L, Chen G, Sun Y-F, Peng L-Y, Chen Y-H. Dynamin-2 mediates heart failure by modulating Ca 2 +-dependent cardiomyocyte apoptosis ☆. 2013.

43. Gemberling M, Karra R, Dickson AL, Poss KD. Nrg1 is an injury-induced cardiomyocyte mitogen for the endogenous heart regeneration program in zebrafish. Elife 2015;4.

44. Egeblad M, Mortensen OH, Kempen LCLT Van, Jäättelä M. BIBX1382BS, but not AG1478 or PD153035, inhibits the ErbB kinases at different concentrations in intact cells. Biochem Biophys Res Commun 2001;281:25–31.

45. Xia W, Mullin RJ, Keith BR, Liu LH, Ma H, Rusnak DW, Owens G, Alligood KJ, Spector NL. Anti-tumor activity of GW572016: a dual tyrosine kinase inhibitor blocks EGF activation of EGFR/erbB2 and downstream Erk1/2 and AKT pathways. Oncogene 2002 2141 2002;21:6255–6263.

46. Anderson NG, Ahmad T, Chan K, Dobson R, Bundred NJ. ZD1839 (Iressa), a novel epidermal growth factor receptor (EGFR) tyrosine kinase inhibitor, potently inhibits the growth of EGFR-positive cancer cell lines with or without erbB2 overexpression. Int J Cancer 2001;94:774–782.

47. Keulenaer GW De, Feyen E, Dugaucquier L, Shakeri H, Shchendrygina A, Belenkov YN, Brink M, Vermeulen Z, Segers VFM. Mechanisms of the Multitasking Endothelial Protein NRG-1 as a Compensatory Factor during Chronic Heart Failure. Circ Hear Fail 2019;12:6288.

48. Reuter S, Soonpaa MH, Firulli AB, Chang AN, Field LJ. Recombinant neuregulin 1 does not activate cardiomyocyte DNA synthesis in normal or infarcted adult mice. PLoS One 2014;9.

49. Reiss K, Cheng W, Ferber A, Kajstura J, Li P, Li B, Olivetti G, Homcy CJ, Baserga R, Anversa P. Overexpression of insulin-like growth factor-1 in the heart is coupled with myocyte proliferation in transgenic mice. Proc Natl Acad Sci U S A 1996;93:8630–8635.

50. Gallo S, Vitacolonna A, Bonzano A, Comoglio P, Crepaldi T. ERK: A Key Player in the Pathophysiology of Cardiac Hypertrophy. Int J Mol Sci 2019;20.

51. Sawyer DB, Zuppinger C, Miller TA, Eppenberger HM, Suter TM. Modulation of Anthracycline-Induced Myofibrillar Disarray in Rat Ventricular Myocytes by Neuregulin-1β and Anti-erbB2. Circulation 2002;105:1551–1554.

52. Kehat I, Davis J, Tiburcy M, Accornero F, Saba-El-Leil MK, Maillet M, York AJ, Lorenz JN, Zimmermann WH, Meloche S, Molkentin JD. ERK1/2 regulate the balance between eccentric and concentric cardiac growth. Circ Res 2011;108:176–183.

53. Peng Y, Zhou B, Wang Y, Chen Y, Li H, Song Y, Zhang L, Rao L. Association between polymorphisms in the signal transducer and activator of transcription and dilated cardiomyopathy in the Chinese Han population. Mol Cell Biochem 2012;360:197–203.

54. Kimura A, Ishida Y, Furuta M, Nosaka M, Kuninaka Y, Taruya A, Mukaida N, Kondo T. Protective Roles of Interferon-γ in Cardiac Hypertrophy Induced by Sustained Pressure Overload. J Am Hear Assoc Cardiovasc Cerebrovasc Dis 2018;7.

55. Rohrbach S, Niemann B, Silber RE, Holtz J. Neuregulin receptors erbB2 and erbB4in failing human myocardium. Depressed expression and attenuated activation. Basic Res Cardiol 2005;100:240–249.

56. Rohrbach S, Yan X, Weinberg EO, Hasan F, Bartunek J, Marchionni MA, Lorell BH. Neuregulin in Cardiac Hypertrophy in Rats With Aortic Stenosis. Circulation 1999;100:407–412.

57. Chen H, Jing XY, Shen YJ, Wang TL, Ou C, Lu SF, Cai Y, Li Q, Chen X, Ding YJ, Yu XC, Zhu BM. Stat5-dependent cardioprotection in late remote ischaemia preconditioning. Cardiovasc Res 2018;114:679–689.

58. Wu Q, Wang T, Chen S, Zhou Q, Li H, Hu N, Feng Y, Dong N, Yao S, Xia Z. Cardiac protective effects of remote ischaemic preconditioning in children undergoing tetralogy of fallot repair surgery: a randomized controlled trial. Eur Heart J 2017;105:151–154.

59. Heusch G, Musiolik J, Kottenberg E, Peters J, Jakob H, Thielmann M. STAT5 Activation and Cardioprotection by Remote Ischemic Preconditioning in HumansNovelty and Significance. Circ Res 2012;110.

60. Mendes-Ferreira P, Keulenaer GW De, Leite-Moreira AF, Brás-Silva C. Therapeutic potential of neuregulin-1 in cardiovascular disease. Drug Discov Today 2013;18:836–842.

61. Odiete O, Hill MF, Sawyer DB. Neuregulin in Cardiovascular Development and Disease. Circ Res 2012;111:1376.

62. Yu CL, Jin YJ, Burakoff SJ. Cytosolic Tyrosine Dephosphorylation of STAT5: POTENTIAL ROLE OF SHP-2 IN STAT5 REGULATION. J Biol Chem 2000;275:599–604.

63. Huang CY, Lin YC, Hsiao WY, Liao FH, Huang PY, Tan TH. DUSP4 deficiency enhances CD25 expression and CD4+ T-cell proliferation without impeding T-cell development. Eur J Immunol 2012;42:476–488.

64. Rigacci S, Guidotti V, Parri M, Berti A. Articles Modulation of STAT5 Interaction with LMW-PTP during Early Megakaryocyte Differentiation. 2008.

65. Nakajima H, Brindle PK, Handa M, Ihle JN. Functional interaction of STAT5 and nuclear receptor co-repressor SMRT: implications in negative regulation of STAT5-dependent transcription. EMBO J 2001;20:6836–6844.

66. Sefat-E-Khuda, Yoshida M, Xing Y, Shimasaki T, Takeya M, Kuwahara K, Sakaguchi N. The Sac3 Homologue shd1 Is Involved in Mitotic Progression in Mammalian Cells. J Biol Chem 2004;279:46182–46190.

67. Martens N, Uzan G, Wery M, Hooghe R, Hooghe-Peters EL, Gertler A. Suppressor of cytokine signaling 7 inhibits prolactin, growth hormone, and leptin signaling by interacting with STAT5 or STAT3 and attenuating their nuclear translocation. J Biol Chem 2005;280:13817–13823.

68. Liu X, Gu X, Li Z, Li X, Li H, Chang J, Chen P, Jin J, Xi B, Chen D, Lai D, Graham RM, Zhou M. Neuregulin-1/erbB-Activation Improves Cardiac Function and Survival in Models of Ischemic, Dilated, and Viral Cardiomyopathy. J Am Coll Cardiol 2006;48:1438–1447.

69. Komamura K, Hanatani A, Ishibashi-Ueda H, Higashi M, Taguchi A, Kangawa K, Nakatani T, Miyatake K. Central Bringing Excellence in Open Access Recombinant Insulin-like Growth Factor-1 Improves Cardiac Function and Symptoms in the Patients on the Waiting List for Heart Transplantation with End-stage Dilated Cardiomyopathy. 2016.

70. Welch S, Plank D, Witt S, Glascock B, Schaefer E, Chimenti S, Andreoli AM, Limana F, Leri A, Kajstura J, Anversa P, Sussman MA. Cardiac-specific IGF-1 expression attenuates dilated cardiomyopathy in tropomodulin-overexpressing transgenic mice. Circ Res 2002;90:641–648.

71. Jie B, Zhang X, Wu X, Xin Y, Liu Y, Guo Y. Neuregulin-1 suppresses cardiomyocyte apoptosis by activating PI3K/Akt and inhibiting mitochondrial permeability transition pore. Mol Cell Biochem 2012;370:35–43.

72. Cohen JE, Purcell BP, MacArthur JW, Mu A, Shudo Y, Patel JB, Brusalis CM, Trubelja A, Fairman AS, Edwards BB, Davis MS, Hung G, Hiesinger W, Atluri P, Margulies KB, Burdick JA, Woo YJ. A bioengineered hydrogel system enables targeted and sustained intramyocardial delivery of neuregulin, activating the cardiomyocyte cell cycle and enhancing ventricular function in a murine model of ischemic cardiomyopathy. Circ Hear Fail 2014;7:619–626.

73. Rohrbach S, Muller-Werdan U, Werdan K, Koch S, Gellerich NF, Holtz J. Apoptosis-modulating interaction of the neuregulin/erbB pathway with antracyclines in regulating Bcl-xS and Bcl-xL in cardiomyocytes. J Mol Cell Cardiol 2005;38:485–493.

